# Stimulus-dependent functional network topology in mouse visual cortex

**DOI:** 10.1101/2023.07.03.547364

**Authors:** Disheng Tang, Joel Zylberberg, Xiaoxuan Jia, Hannah Choi

## Abstract

Information is processed by networks of neurons in the brain. On the timescale of sensory processing, those neuronal networks have relatively fixed anatomical connectivity, while functional connectivity, which defines the interactions between neurons, can vary depending on the ongoing activity of the neurons within the network. We thus hypothesized that different types of stimuli, which drive different neuronal activities in the network, could lead those networks to display stimulus-dependent functional connectivity patterns. To test this hypothesis, we analyzed electrophysiological data from the Allen Brain Observatory, which utilized Neuropixels probes to simultaneously record stimulus-evoked activity from hundreds of neurons across 6 different regions of mouse visual cortex. The recordings had single-cell resolution and high temporal fidelity, enabling us to determine fine-scale functional connectivity. Comparing the functional connectivity patterns observed when different stimuli were presented to the mice, we made several nontrivial observations. First, while the frequencies of different connectivity motifs (i.e., the patterns of connectivity between triplets of neurons) were preserved across stimuli, the identities of the neurons within those motifs changed. This means that functional connectivity dynamically changes along with the input stimulus, but does so in a way that preserves the motif frequencies. Secondly, we found that the degree to which functional modules are contained within a single brain region (as opposed to being distributed between regions) increases with increasing stimulus complexity. This suggests a mechanism for how the brain could dynamically alter its computations based on its inputs. Altogether, our work reveals unexpected stimulus-dependence to the way groups of neurons interact to process incoming sensory information.

## 1 Introduction

Visual information is processed by networks of neurons spanning multiple regions of the neocortex. The interactions between these neurons determine the sensory information extracted by the brain and used to guide behavior. For this reason, much prior work has investigated properties of the networks that define the interactions between neurons in visual cortex. For example, some work has focused on the patterns of anatomical connectivity between individual neurons [1, 2, 3, 4], or between larger voxels of cortical tissue [5, 6, 7, 8]. At the same time, functional connectivity networks – which describe the interactions between neurons – can differ substantially from anatomical networks [9, 10, 11]. Notably, while anatomical connectivity is relatively fixed on the timescale of sensory processing, functional connectivity can vary as the neurons within the network adapt quickly to different stimuli [12]. This motivated us to ask whether and how different stimuli might engage different functional networks with single-neuron resolution within the visual cortex. Despite the clear importance of this question for understanding visual processing, and the substantial literature on functional and anatomical neural network structures (reviewed below), we are unaware of any prior work that addressed how the topological structure of functional connectivity networks between individual neurons spanning multiple regions varies as the stimulus changes. To fill this knowledge gap, we applied network analyses to simultaneous recordings from hundreds of neurons in mouse visual cortex. Our results indicate a surprising degree of stimulus-dependence to the topological structure of functional networks between individual neurons in visual cortex.

Previous work investigated anatomical connectivity between cortical neurons and regions using electron microscopy [13, 4], paired intracellular electrophysiology recordings [1, 2], viral tracing [5, 14], and diffusion tensor imaging [15]. These studies revealed many interesting features of anatomical neuronal connectivity networks, like their modular organization and small-worldness [14, 16, 5, 17], and their hierarchical structure [14]. While anatomical connectivity (e.g., synaptic connections between neurons) remains relatively static over the timescale of processing visual inputs, functional connectivity can be much more dynamic, thus motivating efforts to understand the relation between functional and anatomical connectivity [18, 19, 11, 7, 20]. These efforts are complicated by the fact that different types of stimuli lead to different dynamical patterns of neural activity and to different degrees of correlation between neurons [21, 22, 23, 24, 25]. Because functional connectivity depends on these properties – e.g., on the time-lagged correlation between the activities of neuron pairs [26, 27] – the functional connectivity can depend on the stimulus presented in the experiment.

Despite this potential complication, stimulus-and task-related functional connectivity patterns obtained at a coarse scale using non-invasive functional magnetic resonance imaging (fMRI) have been reported to resemble resting-state functional connectivity patterns [28, 29, 30], while resting-state connectivity in turn resembles anatomical connectivity patterns [31]. In other reports – again, derived from fMRI experiments – stimulus-evoked functional interactions were found to vary with tasks or cognitive states [32, 33, 34, 35, 36]. These fMRI studies raised the important question of whether and how the functional connectivity of the underlying neuronal networks (i.e., at a finer single-neuron scale) might change with stimulus or task conditions.

Functional connectivity at this finer scale is less well-studied due to challenges in simultaneous recordings from large populations of neurons with high spatial and temporal resolution. Despite these limitations, prior work has shown that functional connectivity: 1) is much more stimulus-dependent for high-frequency oscillatory activity than for low-frequency [37]; 2) varies by cell type with the cortex [38]; 3) depends on the contrast of a visual stimulus [39]; and 4) reflects the existence of two main groups of neurons, one whose activities follow those of the rest of the population, and one whose activities do not [24]. While these studies have revealed much about the stimulus-dependence of functional networks at single-neuron resolution, they have not included detailed analyses of networks spanning multiple brain regions. On the other hand, the previous reports of network analysis applied to single-neuron resolution functional connectivity networks, focused on responses mainly to drifting grating stimulus with spontaneous activity as a baseline comparison [26, 27], thus precluding an assessment of stimulus-dependent network structure. Therefore, it is still unclear whether and how the topological organization of these functional networks (either within a brain region, or spanning multiple regions) depends on stimulus properties or other context-defining variables [40].

To fill this gap, we used network analysis methods (similar to those of [26, 27]) to analyze the functional connectivity networks measured in response to 6 different types of stimuli, of varying degrees of complexity, ranging from full-field flashes up to natural movies. These networks were obtained from the simultaneously recorded activities of hundreds of neurons in 6 different cortical regions with implanted Neuropixels probes [26]. Thus, we were able to identify functional networks for each stimulus type, which spanned multiple brain regions. Note that to focus on between-stimulus analyses, we constructed one network based on all conditions for each stimulus, hence the functional networks embody total correlations rather than signal or noise correlations. By studying the structures of these networks and how they varied with stimulus type, we identified several surprising features of the functional networks. First, while the distribution of different types of 3-neuron connectivity motifs were quite similar for the different stimuli, the specific identities of the neurons within those motifs depended on the stimulus. This means that the cortical network is dynamically reorganized as the stimulus changes, but does so in a manner that preserves the motif frequencies. This finding points to a potentially fundamental role for these motif distributions in maintaining the function of the cortical networks [41, 42]. Secondly, we identified highly-interacting *modules* [43, 44] and found that these modules were much more localized to a single brain region (as opposed to being distributed between regions) for stimuli with higher complexity, such as natural movies. Our results thus reveal distinct stimulus-dependent topology of cortical functional networks, and imply a key organizational principle underlying that stimulus-dependence: preserved relative motif frequencies.

## 2 Results

To determine whether and how visual cortical functional connectivity networks depend on the stimulus presented to the animal, we analyzed data from Neuropixels probes inserted into six visual regions of mouse cortex (Fig. 1A: V1, LM, RL, AL, PM, AM), which is previously released by Allen Institute [26]. These probes simultaneously recorded neural activity from each of these six regions while the mice were presented with visual stimuli of varying degrees of complexity (Fig. 1B): flashes, drifting gratings, static gratings, natural scenes and movies, and gray screen (approximation for resting state, or spontaneous activity). From the responses to each stimulus, we extracted the directed functional connectivity using cross-correlograms (CCGs) between the spiking responses of pairs of neurons (Fig. 1C). In order to take polysynaptic connections into consideration [45], we examined ‘sharp intervals’ instead of the ‘sharp peaks’ that might be used to identify monosynaptic connections[46, 47, 26]. These sharp intervals were defined to have a short latency and potentially multiple time lags, and were detected by searching for statistically significantly outlying values in the CCG. Identification of bidirectional connections was made possible by limiting lag *τ* to be non-negative, and each significant connection was defined as excitatory or inhibitory depending on the sign of the significantly outlying CCG value (see Fig. 1C and Methods), similar to the definition used in previous work [48]. Intuitively, if the spiking of the source neuron is statistically strongly correlated with the firing or non-firing of the target neuron with a short time lag, then there exists an excitatory or inhibitory functional connection between them.

**Fig. 1.**
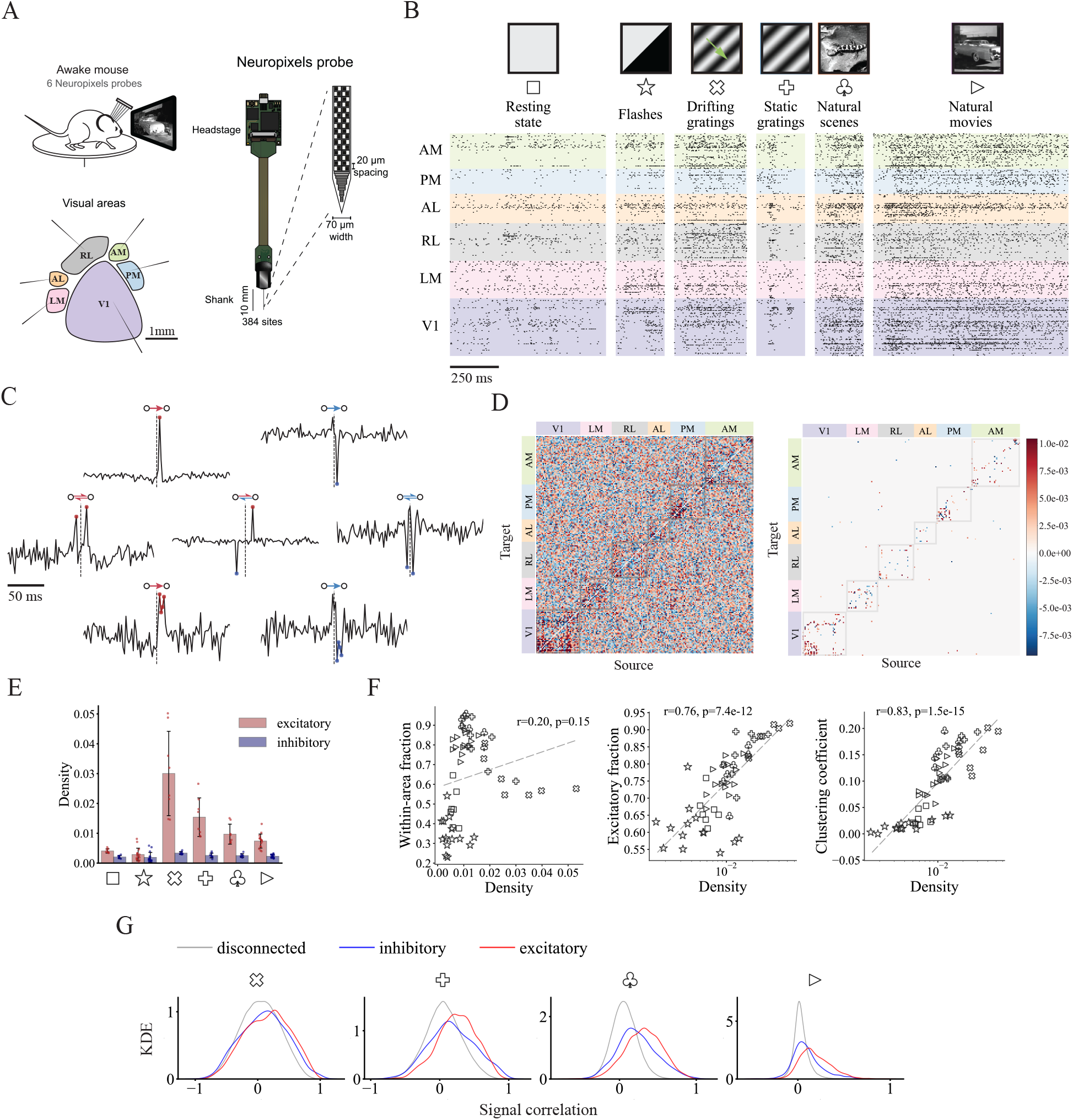
From spike trains to functional connectivity for mouse visual cortex. (A) Schematic of data collection with Neuropixels probes inserted through six visual cortical areas (AM, PM, AL, RL, LM and V1). (B) Example spike trains of 193 units from the visual cortex of a mouse during six different types of stimuli. For brevity, each stimulus is denoted using a unique symbol in all figures. (C) Example CCGs (cross-correlograms) of excitatory (red)/inhibitory (blue), unidirectional/bidirectional and monosynaptic (‘sharp peak’)/polysynaptic (‘sharp intervals’) connections. (D) (left) Example matrix of CCG with units ordered by area during natural movie stimuli. (right) Connectivity matrix with only significant connections (*Z >* 4). (E) Density of excitatory and inhibitory connections during all visual stimuli. Density is defined as the number of connections normalized by total possible number of connections. (F) Fraction of within-area connections, fraction of excitatory connections and clustering coefficient against network density. Each visual stimulus is characterized by a symbol, consistent with (B). (G) Kernel density estimation (KDE) of signal correlation distributions for disconnected neuron pairs and pairs with inhibitory/excitatory connections during presentations of four types of visual stimuli.

To obtain a comprehensive understanding of the stimulus-dependent structure of the functional connectivity networks (Fig. 1D), we conducted network analyses at multiple topological scales, ranging from the properties of pairwise connections to the local connectivity patterns of third-order functional motifs, up to larger-scale functional modules. Our control analysis on running speed (not shown) showed that our subsequent observations are indeed determined by the stimulus and not by locomotion.

### 2.1 Stimulus dependency of functional connectivity networks

We first investigated overall patterns of functional connections between neurons across stimulus types by comparing the functional connectivity matrices. We found there are some common network features observed across stimulus types. Specifically, the functional networks observed during all visual stimuli exhibited heavy-tailed degree distributions (Supplementary Fig. 1B). Networks with this property are known to be robust to random failures [49], however, they are more vulnerable to targeted attacks on hub neurons which could lead to reduced network efficiency as observed in Alzheimer’s patients [50, 51].

While functional networks show some shared characteristics like heavy-tailed degree distributions across stimuli, we also observed network properties vary with stimulus complexity. We found that natural stimuli (natural scenes and movies) tended to evoke fewer functional connections than grating stimuli (both static and drifting gratings) while full field flashes drive the least correlated neural activities, on the same level as resting state activity (Fig. 1E). These findings are consistent with previous reports that natural stimuli decorrelate neurons in primary visual cortex (V1) [52, 53]. While these previous works focus on V1, our results suggest that decorrelation by natural stimuli is a general property of cortical circuits: it is found in higher visual cortical areas as well.

The differences in network density mainly originate from differences in excitatory connections (Fig. 1E), which results in the strong correlation between the fraction of excitatory connections and the network density (Fig. 1F, middle). Even though natural stimuli do not evoke the densest functional networks, the fraction of within-area connections is largest for static gratings and natural stimuli (Fig. 1F, left, and Supplementary Fig. 1A; *p <* 10*^−^*^3^, Kolmogorov-Smirnov test). This is closely related to the stimulus-dependent differences in modular network structure, which we analyzed in more details later in this paper.

To determine how the stimulus-dependence of the network density affects the network’s topological structure, we measured the tendency for triplets of neurons to form closed triangles (e.g., three-neuron motifs 6,9-13 in Fig. 2B). This tendency is quantified by the clustering coefficient, and we found that it increases with increasing network density regardless of stimulus type (Fig. 1F, right).

**Fig. 2.**
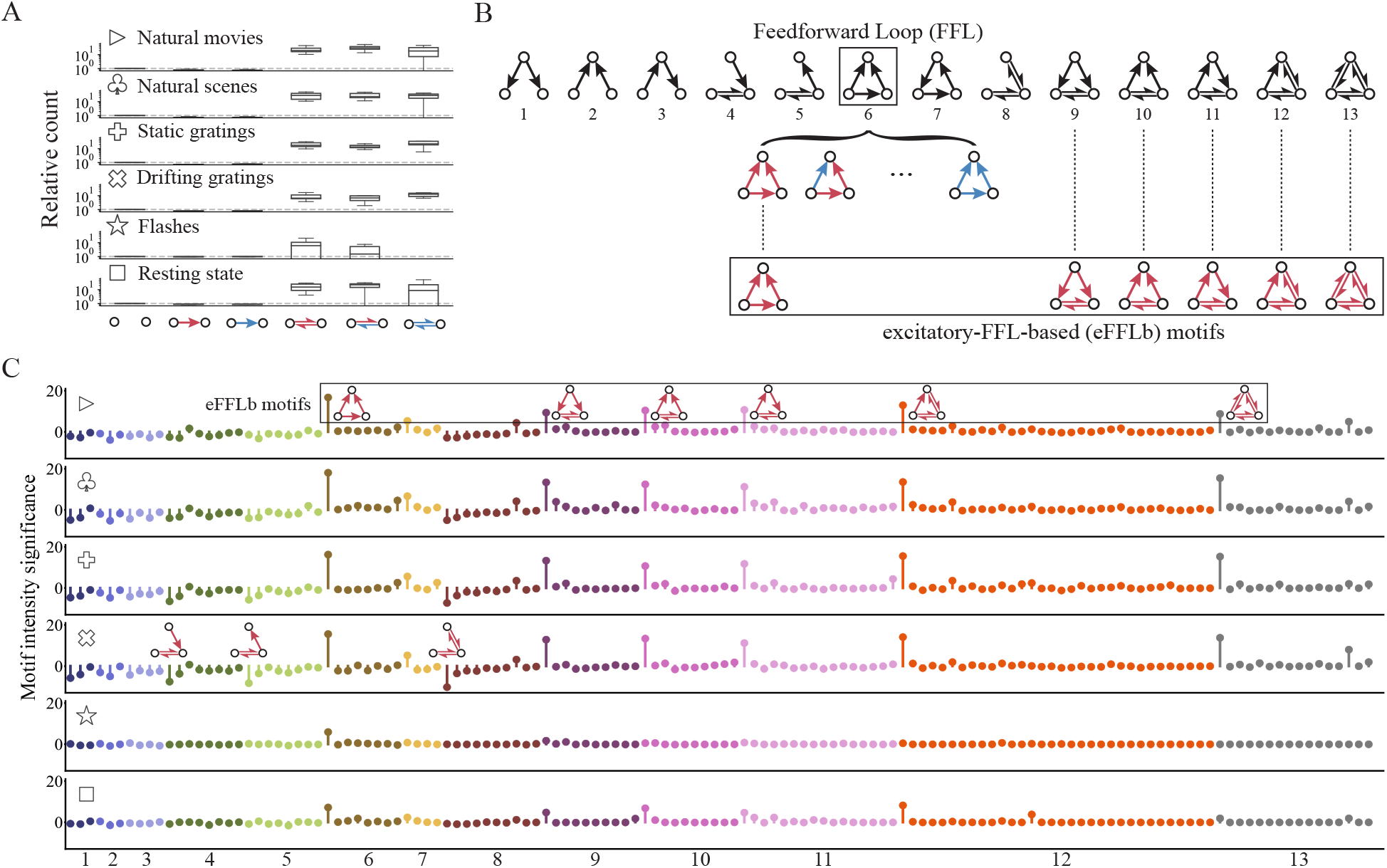
Highly preserved local structure during different types of visual stimuli. (A) Relative count of signed neuron pairs with respect to reference model (Erdős-Rényi model). All three types of bidirectional pairs are considerably over-represented. (B) 13 types of motifs without edge signs. Examples of signed motifs based on Feedforward Loop structure (motif ID = 6) are shown. Excitatory-FFL-based (eFFLb) motifs are defined as motifs with at least one FFL structure consisting of all excitatory connections. Note that eFFLb motifs can be classified into four types based on the number of mutual connections: zero (ID = 6), one (ID = 9, 10, 11), two (ID = 12) and three (ID = 13). In what follows, we use the prefix “e” or “i” to denote signed motif types with only excitatory connections or only inhibitory connections, respectively. (C) Motif intensity significance sequences of all signed motifs (signed three-neuron subnetworks). Each row shows results for a certain stimulus type and each color corresponds to a certain unsigned motif structure. Motif intensity significance for each signed motif is obtained through its intensity Z score of the empirical network with Signed-pair-preserving model as the reference. The overall sequences are remarkably similar across visual stimulus classes, with six types of over-represented motifs (eFFLb motifs) and three under-represented motifs preserved. It is worth noting that the above-mentioned nine significant motifs only contain excitatory connections.

Motivated by previous work showing that neurons with similar preferences tend to connect with each other [2, 4, 54], we compared the tuning similarity of neuronal pairs connected with excitatory and inhibitory connections. To perform this comparison, we computed the kernel density estimation (KDE) for signal correlation during presentation of four visual stimulus types. Signal correlation is defined as the correlation between average responses of neurons to different stimulus conditions which is used to test whether two neurons have similar tuning curves [21, 55]. We computed these signal correlation separately for pairs with excitatory connections, inhibitory connections, and those with no connections (Fig. 1G). For natural movies, we regarded each frame as a different stimulus condition when computing the signal correlation [2]. Since there are only two conditions for flashes (dark or light), the signal correlation of either 1 or -1 could be trivial and thus is not considered in this analysis. Similarly, the signal correlation is ill-defined for the blank gray screen stimulus, and thus it was also omitted from this analysis.

For all visual stimuli, the signal correlations for connected neuron pairs tended to be larger than for disconnected pairs, which had distributions centered around zero (Fig. 1G; *p <* 10*^−^*^144^, rank-sum test). Additionally, neurons with excitatory connections tended to have higher signal correlations than did pairs with inhibitory connections (Fig. 1G; *p <* 10*^−^*^3^, rank-sum test).

In agreement with the previous findings that neurons close in space or sharing similar tuning curves are more likely to have synaptic connections [54, 2], we found the probability of functional connections decreases with increasing distance and increases with their increasing signal correlation (Supplementary Fig. 2A,B; Cochran-Armitage test). In addition, the probability of a functional connection being excitatory/inhibitory significantly increased/decreased with signal correlation during all visual stimuli (Supplementary Fig. 2C,D; Cochran-Armitage test), indicating that even though neurons with similar preferences generally tend to be connected, the sign of the connection depends on the extent of their tuning similarity.

Collectively, these analyses show that network density, fraction of connections that are within a brain region (as opposed to between regions), clustering coefficient, and the distribution of signal correlation, depend systematically on the stimulus type.

### 2.2 Stimulus dependency of functional connectivity motifs

Having observed stimulus-dependency of the general network properties, we next turned our attention to the properties of the functional motifs. Specifically, we investigated two-and three-node motifs in the functional connectivity network. Similar to anatomically-defined structural motifs which form fundamental building blocks of neural circuits [56, 1, 57], functional motifs, which are defined by correlated neural activities, represent elementary information processing components of a functional network [58].

To understand the distribution of two-and three-neuron motifs in each functional network, we adopted the intensity method for functional motif detection. This method computes the Z-score of the intensity for a given motif by comparing the motif frequency in the empirical network and in a randomly-generated surrogate network [59]. This method thus identifies how much more (or less) prevalent the motif is in the real network than would be expected by chance in a reference network randomly shuffled with certain preserved properties (e.g., density, degree distribution, etc).

Note that, to characterize pairwise signed connections (i.e. signed two-neuron motifs), it is necessary to preserve the edge sign distribution when generating density-matched randomized surrogate networks[60]. For this reason, we included edge signs in the pair-preserving model [56] that preserves the distribution of (n-1)-neuron motifs and used the resultant Signed-pair-preserving model to generate the surrogate network with the preserved signed (n-1)-neuron motifs for comparing the motif frequency between the true network and the randomly-generated one.

As for two-neuron motifs, we found that bidirectional signed functional connections are much more frequent than would be expected by chance in the random network (Fig. 2A). This recapitulates the structural observations that bidirectional synaptic connections are highly over-represented in cortex relative to density-matched random networks [1].

We further studied distributions of three-neuron motifs in functional networks. There are 13 types of connected three-neuron motifs (Fig. 2B), of which the Feedforward Loop (FFL) is arguably the most studied type due to its ubiquitous nature in empirical networks such as gene systems and neuronal networks [61]. The lollipop plot in Fig. 2C shows the intensity Z score obtained using the above intensity method for all types of signed three-neuron motifs ordered by their corresponding unsigned motif types. The colors in the plot represent each unsigned type (see Supplementary Fig. 4 for the full set of signed connectivity patterns). Interestingly, the most salient motifs were the same for the different visual stimuli. Specifically, the top six over-represented motifs are the same for different stimuli being tested here (Fig. 2C, motif ID = e6, e9, e10, e11, e12, e13). In addition to these significantly over-represented motifs, the same three types of motifs were significantly under-represented for all stimuli (ID = e4, e5, e8). Interestingly, all 6 over-represented motifs contain at least one excitatory FFL (eFFL) structure (ID = e6, Fig. 2C) with the only difference among them being the number of mutual connections that accompany the eFFL structure. The last five of these over-represented motifs were studied without edge signs in previous works as ‘mixed-feedforward-feedback loops’ and have been found to be correlated to memory, as well as acceleration and delay of response [62]. Here we denote all six of the over-represented signed motifs as excitatory-feedforward-loop-based (eFFLb) motifs.

In addition to the three highly under-represented motifs (ID = e4, e5, e8), the set of under-represented motif patterns consists largely of ‘unclosed’ eFFLb motifs, suggesting that neurons tend to form pairwise-connected triplets. This under-representation of unclosed motifs appears to be more pronounced with higher network density. Interestingly, the motifs’ average absolute intensity Z scores (deviation from the frequency expected by chance) increases significantly with increasing network density (Fig. 3A). With more connections, the empirical functional network deviates more strongly from randomized surrogate networks, highlighting the fundamental non-randomness of local functional connectivity.

**Fig. 3.**
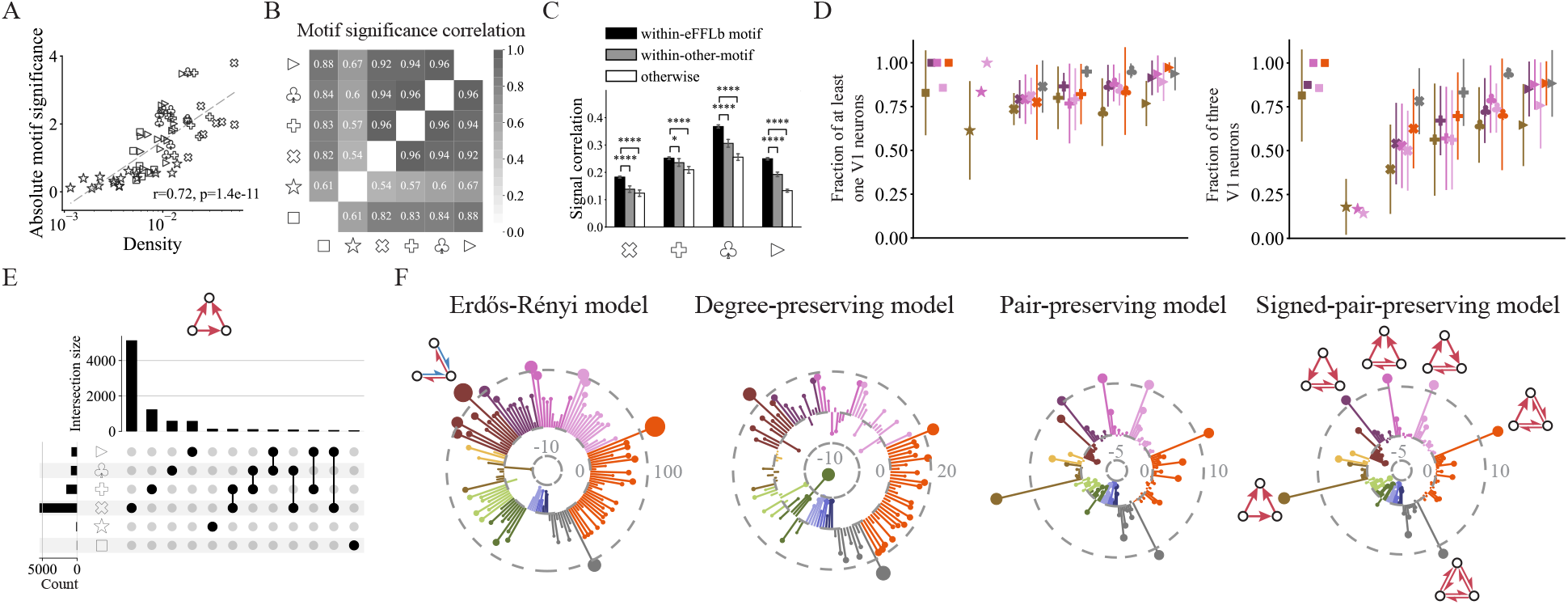
Same motifs and similar patterns are organized from different neurons. (A) Average absolute motif significance (absolute Z score of intensity) across all signed motifs against network density. Similar to within-area fraction and clustering coefficient (Fig. 1F), there is also a logarithmic relationship between motif significance and density. (B) Pairwise correlation of normalized motif intensity distribution for six visual stimuli. Extremely high correlation proves the similar motif presence during different types of stimuli. (C) Signal correlation during four visual stimuli (except for resting state) for within-eFFLb-motif, within-other-motif connections and others. Other over-represented motifs are determined using a significance level of 99%. *p <* 0.05, *p <* 0.0001, rank-sum test. (D) Fraction of motifs with at least one V1 neuron or all three V1 neurons during all visual stimuli for six over-represented excitatory-feedforward-loop-based (eFFLb) motifs. Six colors represent six types of eFFLb motifs, consistent with Fig. 2C. From left to right, motif ID = e6, e9, e10, e11, e12, e13. (E) Intersections of unique motif sets for motif ID = e6 during six types of stimuli. Horizontal bar plot shows the size of each intersection set while vertical bar plot displays the number of signed motif ID = e6 for each type of stimulus. A unique motif is defined as a certain signed motif with three specific neurons, and intersections with less than 20 elements are removed for brevity (see Supplementary Fig. 4 for the complete results). A large number of unique motifs appear only during one type of stimuli, demonstrating that even though functional motifs are preserved across visual stimuli, component neurons are changing. (F) Multiple motif intensity significance sequences were obtained through four different reference models for natural movies as the representative stimulus type: Erdős-Rényi model, Degree-preserving model, Pair-preserving model and Signed-pair-preserving model with an increasing number of preserved network properties (Methods). Colors are consistent with (D) and Fig. 2C, and connectivity pattern is shown only once for each type of most significant signed motif for brevity. Empirical functional networks are progressively more similar to surrogate networks from left to right.

Note that it is important to examine the whole significance distribution of motifs instead of focusing on merely the most striking ones. Varying the threshold on significant motifs naturally changes their total count, but importantly, it does not substantially change the relationship between the functional connectivity patterns observed for the different stimuli (Supplementary Fig. 5B). Our control analysis has led us to the conclusion that the presence of non-random local topology of functional networks is contingent upon the density of the network(Fig. 3A,B), which is, in turn, modulated by sensory input (Fig. 1E). Meanwhile, the same sets of two-and three-neuron connectivity motifs are over-and under-represented for all 6 visual stimuli.

Our analysis of the connectivity motifs relies on comparing motif frequencies in the observed network to those reference networks that are similar in some way to the observed network but are otherwise randomized. Notably, there are many different ways to define random networks by preserving certain network properties. For example, the commonly used Erdős-Rényi reference network preserves the network density. Other reference models can preserve the degree distribution, the neuron pair distribution, or the signed neuron pair distribution (see Methods). To understand how our motif analysis depends on the choice of reference model, we computed the three-neuron motif intensities for the natural movies stimulus for 4 different reference models with increasingly more preserved network properties (from left to right in Fig. 3F). As a result, the overall significance level for all motif types roughly decreases in the same order, and the most strictly conserved reference model, the signed pair-preserving reference model, is better in identifying the small subset of motifs for which the observed network is most truly non-random. Therefore, we use the signed pair-preserving reference model as our default reference model throughout this study.

To test the robustness of our results, we separately calculated the motif distribution patterns on distinct halves of the trials from our natural scene stimulus data. The high consistency between the motif distributions on the two data splits (Fig.S11) suggests that noise within the dataset is unlikely to be a key factor in our results. Moreover, eFFLb motifs remain to be the most salient motifs across different stimuli on various significance levels (Fig.S12). Taken together, eFFLb motifs seem to be reliably the most significant patterns among three-neuron subgraphs.

### 2.3 Properties of over-represented three-neuron motifs

While the three-neuron functional connectivity motifs that were most over-or under-represented were the same for all stimuli (Fig. 3B), those motifs were composed of different neurons for different stimuli (Fig. 3D,E). Specifically, while most eFFLb motifs contained at least one neuron from V1, the fraction containing all 3 neurons within V1 varied substantially as the stimulus changed. (Fig. 3D).

To further quantify the extent to which the same neurons constitute eFFLb motifs across stimuli, we computed the number of eFFLb motifs sharing exactly the same neurons for all pairs of stimuli (intersection sizes), as shown in Fig. 3E for one example over-represented eFFLb motif (ID=e6; results for all of the over-represented eFFLb motifs are shown in Supplementary Fig. 6). We also compared motif overlapping detected using half the trials of the same stimulus and using different stimuli (Supplementary Fig. 11). Notably, eFFLb motifs were more likely to be composed of the same neurons during the same stimulus than for different stimuli, indicating that the identities of the neurons within the over-represented motifs change as the stimulus changes.

Taken together, these results and those in Fig. 2C, indicate that, as the stimulus changes, the same three-neuron motifs are over-or under-represented in the cortical networks, but the identities of the neurons within those motifs change. This suggests that these specific motifs might have strong functional importance for the cortical microcircuit because even as different stimuli dynamically alter the functional connectivity, they do so in a way that preserves these motif patterns.

The fact that the eFFLb motifs were over-represented for all stimuli even with different constituent neurons suggested that they might play an important role for the cortical microcircuit. To further investigate this question, we analyzed the tuning similarity of these motifs. We found the signal correlation between pairs of connected neurons within the same eFFLb motif were significantly higher than those not within the same eFFLb motif (Fig. 3C; *p <* 0.05, *p <* 0.0001, Student’s t-test), and neurons within the same eFFLb motif are spatially closer to each other than otherwise (Supplementary Fig. 5A, right; *p <* 10*^−^*^4^, Student’s t-test). Furthermore, neuron pairs within eFFLb motifs show stronger functional connection strengths and higher CCG Z-scores than other connected neuron pairs (Supplementary Fig. 5A; *p <* 10*^−^*^4^, Student’s t-test). Thus, the eFFLb motifs tend to consist of functionally-similar neurons. These observations are consistent with previous reports of synaptic connectivity patterns in visual cortex [1].

Overall, these analyses indicate that neuron pairs within the over-represented eFFLb motifs tend to be spatially near each other, and to have higher functional similarity compared to other pairs. Coupled with the fact that these eFFLb motifs are preserved across stimuli, this highlights the potential functional importance of these motifs within the cortical microcircuit.

### 2.4 Spatial and functional organization of network modules depends on the stimulus

Having identified the fundamental motif structures as computational building blocks regardless of stimulus types, we next asked how the topology of larger groups of neurons, or modules, depends on stimulus properties. Those *modules* [63, 64] are thought to impart added robustness [65], efficiency [66], and functional specialization [44] to networks. We thus sought to identify modules within our networks, and to determine how their properties depend on the stimulus presented to the animal.

To achieve this goal, we revised the Louvain method [67] to optimize the Modularity estimation from previous work [68, 69] so as to take into account the signs of the connections in our networks: this modified Louvain method searched for sets of modules with many excitatory connections inside the same module and inhibitory connections between different modules (see Methods). Thus, the method identifies sets of modules whose neurons are internally correlated and externally anti-or un-correlated. This greedy optimization yields the groupings of neurons into modules by maximizing the score of modified Modularity (see Methods). For comparison, we also identified modules with the original Modularity algorithm that does not take into account the edge signs [65]. Although our adapted method revealed results qualitatively similar to the original one (Supplementary Fig. 10), the identified module size using our method is relatively smaller, suggesting a finer scale module detection with our method.

Unless otherwise stated, in the rest of this paper Modularity means the modified Modularity for signed module detection. By maximizing the two-dimensional Modularity difference map whose dimensions correspond to resolution parameters for excitatory and inhibitory links, we determined the optimal resolution parameters that control the scale of identified modular structure so that the empirical network deviates most from the null model [70, 14] (see Methods; Supplementary Fig. 7A). In light of the potential limitations posed by a fixed resolution parameter, we analyzed multi-resolution module partitioning and found consistent results (Supplementary Fig. 7B,C,D).

After identifying the best parameters, we applied our module detection algorithm to the observed functional networks from each stimulus, and compared the results between stimuli. During gratings and natural stimuli, functional networks tend to exhibit stronger modular structures, characterized by larger deviations in Modularity from expectation (Fig. 4A, bottom z-score, 53.94 ± 8.81). On the contrary, the networks obtained from flashes and in the resting state were less modular (6.22 ± 4.44; *p* = 3.4 × 10*^−^*^9^, rank-sum test).

**Fig. 4.**
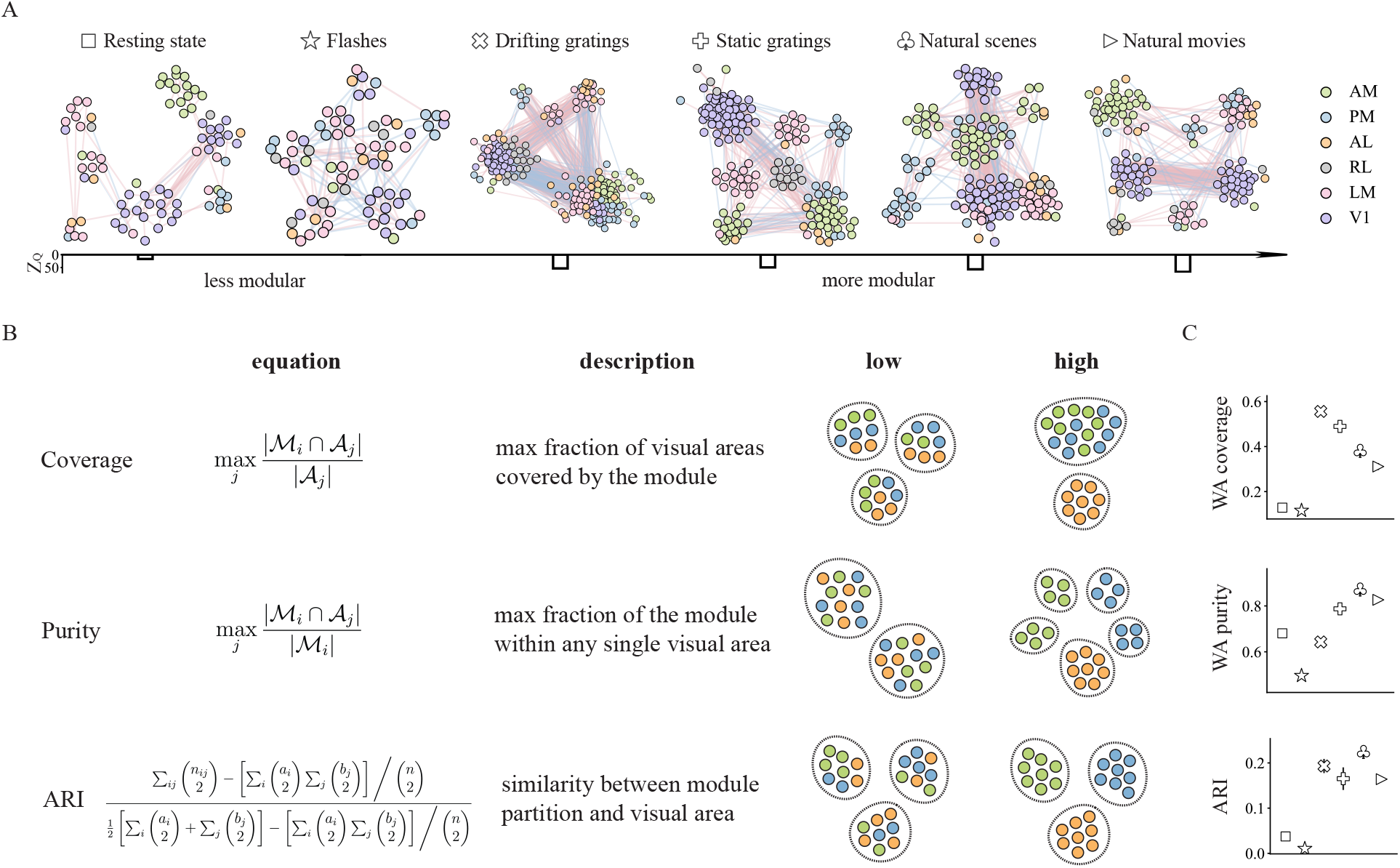
Distinct modular structures during different types of visual stimuli. (A) Topological structure of functional connectivity of a mouse during six types of visual stimuli with neurons colored by area. The color of each connection shows its sign with red denoting excitatory connection and blue representing inhibitory correlation. The community partition is obtained through modified Modularity for signed networks (see Methods). We computed the Z score of Modularity with Signed-pair-preserving model as the reference to show the degree to which functional network has a modular structure. Networks during gratings and natural stimuli show significant modular structure. (B) We used three measures to reveal the modular structure regarding visual area from different perspectives. Coverage and purity are module-level measures, where the former marks the degree to which the module covers any visual area, while the latter measures the degree to which all neurons in the module are from the same visual area. We computed the average coverage and purity weighted by module size to show the overall properties of the whole functional network (see Methods). Adjusted Rand Index (ARI), a network-level measure, was also used to quantify the difference between module partition and visual areal organization. The weighted average (WA) coverage is 0.375 and 1 (ranges from 0 to 1), WA purity is 0.333 and 1 (ranges from 0 to 1) and ARI is -0.03 and 1 (ranges from -0.5 to 1) for the corresponding two toy examples visualizing the ‘low’ and ‘high’ cases for the measure. (C) WA coverage, WA purity and ARI during six visual stimuli. The error bars show the confidence intervals over all mice obtained with non-parametric bootstrap method. In general, there tend to be fewer and larger modules with higher coverage during grating stimuli, whereas we usually find more and smaller modules with higher purity during natural stimuli. As a result, ARI is lower during resting state and flash while higher during grating and natural stimuli.

Anatomically parcellated brain regions are thought to work as natural modules with specialized functions [71, 72, 73, 74]. We thus wanted to understand how our functionally-defined modules relate to the anatomically-defined brain regions. To achieve this goal, we analyzed the identified functional modules to understand the extent to which their spatial organization coincided with the anatomically-defined brain regions, and the extent to which that depended on the stimulus. To do this, we computed three measures from the modules for each stimulus (Fig. 4B, left). First, the *coverage* quantifies the maximum extent to which a given module covers all of the neurons in any single brain region. Second, the *purity* quantifies the maximum extent to which a given module is contained within any single brain region. These two quantities are computed for each module, and the results in Fig. 4 show their weighted average (averaged over modules, weighted by module size). A more detailed module-by-module analysis is presented in Fig. 5, below. Finally, the *adjusted rand index (ARI)* quantifies the similarity between how the modules partition the set of neurons, and how the brain regions partition the set of neurons. Intuitively, these measures revealed the properties of modular structure from different perspectives: high coverage means at least one visual area is covered by the module, high purity means that a module consists of neurons from the same visual area, and high ARI means the overall module partitioning highly resembles the areal organization.

**Fig. 5.**
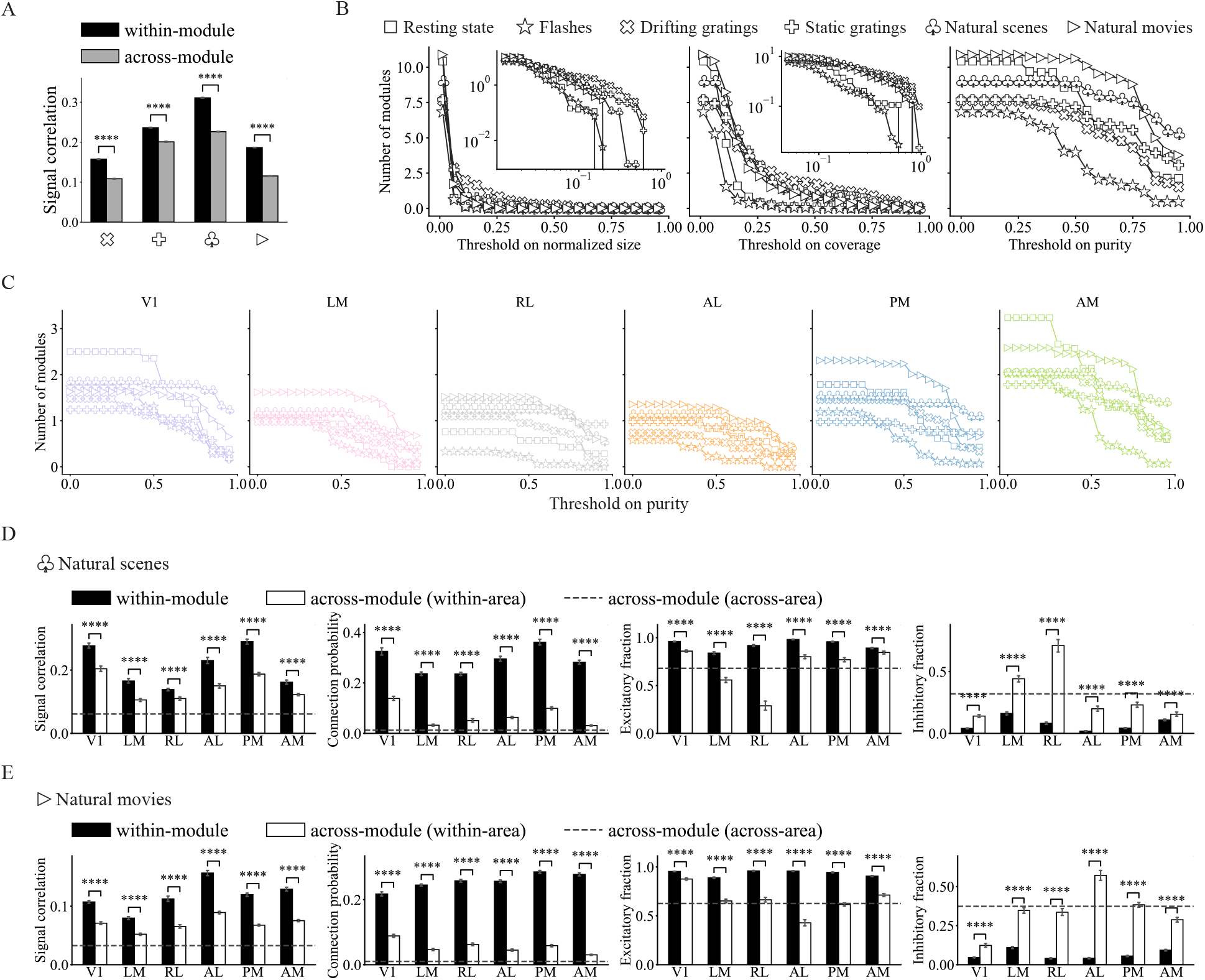
Stronger segregation during natural stimuli. (A) Signal correlation for within-module and across-module connections. *p <* 10*^−^*^4^, Student’s t-test. Connected neurons partitioned into the same module tend to have higher signal correlations than connected neurons from different modules, demonstrating our module partition provides insight into not only the connectivity pattern but also functional similarity to some extent. (B) Number of modules with normalized size, coverage or purity higher than the threshold. Normalized size is the size of module normalized by the total number of neurons in the network, insets show the plots on a log-log scale. (C) Number of modules with purity higher than the threshold for each visual area during all visual stimuli. (D, E) Properties of the modular structure during natural scene and natural movie presentations. We examined four different aspects of the case where neurons from a single visual area are divided into multiple modules (in which they are the dominant area), with signal correlation indicating the functional similarity along with connection probability, excitatory fraction and inhibitory fraction demonstrating the validity of our module partitioning algorithm. ∗ ∗ ∗ ∗ *p <* 10*^−^*^4^, Student’s t-test.

These three measures all show variation in the module organization for different stimuli (Fig. 4C). For natural images and movies, the modules have a higher propensity to cover only a subset of a brain region than for drifting and static grating stimuli. This is reflected by lower coverage scores and higher purity scores for the natural image and movie networks than for the drifting and static grating networks (Fig. 4C; *p <* 10*^−^*^310^, *p <* 10*^−^*^310^, rank-sum test).

It is important to note that with increasing module size, coverage tends to increase while purity tends to decrease, and the module size does depend on the stimulus (Fig. 5B). As a result, it is important to ask whether the variations in coverage and purity with visual stimuli could be explained simply by stimulus dependence of module size. To address this question, we compared module purity and coverage to module size (Supplementary Fig. 8). Consistently across module sizes, the modules in the flashes stimulus and resting state networks had lower coverage than did the networks for the other stimuli. The network from the flashes stimulus also had consistently lower purity.

To further probe the relationships between module sizes and coverage or purity, we analyzed the number of modules obtained for each stimulus that were above a given threshold of module size, threshold of module coverage, or threshold of module purity. Repeating this for many threshold values (Fig. 5B), we found that natural images and movies had the largest numbers of high-purity modules even though their numbers of small modules were not appreciably different from the other stimuli. These findings emphasize that the stimulus-dependent module properties we report in Fig. 4C cannot entirely be attributed to stimulus-dependent module sizes.

In general, the similarity between functional module partitioning and the anatomical areal organization is higher during gratings and natural stimuli and lower during resting state and flashes (*p <* 10*^−^*^310^, rank-sum test), suggesting that functional connectivity tends to be more constrained by the anatomical structure and more spatially compact during complex and natural stimuli. This is reflected in the lower ARI values for the flashes and spontaneous activity (Fig. 4C; *p <* 10*^−^*^310^, rank-sum test).

Consistent with previous work [58, 75], functional motifs seem to be more pronounced in more modular networks (Supplementary Fig. 9 F), suggesting their shared organizational principles. Similar to motifs, we also tested the functional similarity of nodes within and across modules by measuring the signal correlations of connected neuron pairs. Neuron pairs within the same module had higher signal correlations than did neuron pairs in different modules (Fig. 5A), and the probability of any two connected neurons being in the same module also increases with increasing signal correlation (Supplementary Fig. 9A; Cochran-Armitage test). These findings were consistent for the 4 visual stimuli for which the signal correlations are well-defined, and were consistent across brain regions when modules were assigned to the brain region from which most of their neurons came (Fig. 5D,E). These observations emphasize that the modular structure promotes functional specialization [44].

Our analyses of the modular organization of the functional connectivity networks reveal that the modules tend to contain neurons with similar stimulus tuning, and that their spatial organization and alignment with anatomical brain regions depend on the stimulus presented to the animal. This emphasizes that a functional module is not strictly the same as an anatomical brain region: the relationship between these concepts depends on the stimulus-defined context.

## 3 Discussion

We studied the topology of micro-scale functional networks measured with single-neuron spiking activity in the mouse visual cortex. These data were collected while the mice were exposed to different types of visual stimuli, and we separately analyzed the functional networks observed in the responses to each stimulus type. Thus generated functional networks could differ from the underlying anatomical connectivity, and this disparity warrants caution when interpreting the connectivity graphs. However, our science question, concerning stimulus-dependent interactions between neurons cannot be answered with the standard anatomical connection methods. For this reason, we used functional connectivity measures for this study.

We found that functional networks display stimulus-dependent network properties such as varying density, clustering coefficient and fraction of excitatory connections. Furthermore, we provide evidence that the distribution of low-order connectivity patterns (motifs with 2 or 3 nodes) remains stable, characterized by over-representation of a specific group of 3-neuron motifs, eFFLb motifs. This over-representation was preserved across the wide range of stimuli we investigated. Notably, while this motif was over-represented in all cases, the constituent neurons within that motif changed. Finally, we observed that the module-level network architecture depends significantly on the stimulus.

The consistent over-representation of eFFLb motifs suggests that they are key information-processing components of neural circuits. While these motifs were over-represented for all stimuli, the identity and areal distribution of neurons constituting the eFFLb motifs differed between stimuli. Thus it is the three-neuron *patterns* rather than the triadic interactions of specific neurons that are preserved. This observation suggests that an important computational role might arise at the motif level[76, 77, 78, 79, 80], where neurons can dynamically reorganize to form these relevant structures. These local computations, organized by motifs, could remain robust to changes like the loss of individual neurons because other neurons could be recruited into the motifs to replace any that are lost. For this reason, motif-level computational organization could provide substantial robustness to cortical computation.

The abundance of FFL motifs has been observed in numerous types of networks including gene regulatory networks [61], transportation networks [81], engineered networks [56] and neuronal networks[1, 56, 81]. Motif ID = e6 (FFL) is proven to be a sign-sensitive filter that responds only to persistent stimuli in transcriptional regulation networks [61] and multi-input FFL generalization is found to store memory as well as reject transient input fluctuations in neuronal networks [41]. For anatomical networks of neurons, there have been modeling works showing that certain eFFLb motifs (e9, e10) can function as long-term memories of the input, thus playing an important role in most cognitive tasks [62].

Therefore, the eFFLb motifs that we found to be consistently over-represented in cortical functional networks may have important functional roles in cortical computation.

On the global scale, however, more complex visual stimuli tend to drive networks into more modular structures with stronger segregation and stronger agreement between structural and functional parcellation. This suggests that functional modules with more spatially segregated structure could be required in more demanding cognitive tasks. Nevertheless, major functional modules observed when the animal was viewing natural scenes and movies are highly overlapping at all spatial scales with almost the same neurons. These probably arise from the common subtasks required by the visual processing of similar stimuli. One advantage of having shared modular components is that it allows a faster adaptation and possibly a lower switching cost of functional networks to various tasks [75]. This reduction in functional reorganization costs could be especially important given our observation that different neurons are organized into the over-represented eFFLb motifs in the presence of different stimuli.

Anatomical structure has been known to stay relatively stable given different types of sensory inputs[82]. In comparison, functional connectivity changes in such a fast and dramatic manner that some even try to model its temporality within a single trial [83]. There are various reasons for this rapid functional adaptation, which could be a change of task [32, 33, 34, 35], perceptual states [84], visual stimuli [85, 22, 23], etc. However, most prior studies were either restricted to the primary visual cortex, or to voxel-level recordings obtained through fMRI, or both. It thus remains unclear whether the functional interactions on a single-neuron scale across multiple cortical regions are also dynamically adapted to the visual inputs. One major aim of our study was to fill this knowledge gap. By studying interneuronal functional connections, our work could help improve our understanding of neuron-to-neuron connections (e.g., at the synaptic scale). In contrast, studies of voxel-scale functional connectivity based on fMRI data might be less informative about these finer-scale interactions.

One of the major challenges of studying functional connectivity lies in the existence of inhibitory correlations and anti-correlated patterns: there is a lack of strong theoretical tools for analyzing networks with both positive and negative edges [86]. While many works on functional connectivity disregard inhibition and only focus on excitatory connections for simplicity, inhibitory connections play a crucial functional role in visual processing [87, 88]. This highlights the need for a network analysis framework that possesses the ability to handle both positive and negative edges. We address this problem by adopting and modifying motif and module detection methods for signed networks, and used these methods to investigate how (signed) functional networks vary on local and global scales. The inclusion of edge sign in motif analysis enables us to further distinguish motifs, since the same unsigned connectivity could correspond to different functions depending on the edge signs [61]. The definition of functional modules can be subjective regarding whether to keep inhibitory connections inside or between modules. Nonetheless, we showed that even though qualitatively similar conclusions can be drawn without distinguishing functional inhibition from excitation (Supplementary Fig. 11), ignoring edge sign (as in prior works) could lead to a less detailedunderstanding of the exact pattern of functional segregation and specialization.

While our detailed methods thus provide a more comprehensive analysis of how functional connectivity flexibly adapts to the statistics of visual input, we recognize some limitations to our analyses. First, we do not distinguish neurons according to their cell types. This limits our ability to relate our functional connectivity results to the growing literature on microcircuit architectures. In addition, due to incomplete recording, we observe only a subset of the neurons in each brain area. Furthermore, correlation-based network inference can potentially lead to false direct edge identification via high-order connections. These limitations are inherent to neural activity-based construction of functional connectivity, and not just to our study. Nevertheless, we do not believe that these common limitations constitute serious flaws in our analysis. Here we are not trying to find functional networks that topologically resemble the anatomical network (in which case the incomplete recording issue would be quite detrimental). Instead, our focus is on the adaptation of inter-neuronal interaction patterns to different visual stimuli. These interactions can be identified even in incomplete recordings. On the other hand, incomplete observation might explain the presence of nonconforming edges in some of our analyses, such as the presence of both excitatory and inhibitory edges emanating from a single neuron. At first glance, these neurons are at odds with Dale’s principle, which suggests that such bivalent neurons are very uncommon in the neocortex. However, given the incomplete recordings, there could be unobserved inhibitory neurons that mediate the effective inhibitory impact of an excitatory neuron on some other neurons in the circuit.

Additional limitations of our study arise from experimental constraints and the nature of the Neuropixels dataset collected from extracellular electrophysiology probes. Kilosort2 was used to identify spike times and assign spikes to individuals [89], however, no current spike sorting algorithm can ensure a completely accurate assignment of observed spikes to individual neurons. This means some certain nodes in our network could correspond to more than one neuron, or that there could be multiple nodes corresponding to the same neuron [90]. Finally, the limited set of visual stimuli used in our experiments could introduce bias into our analysis since we do not have multiple different sets of stimuli within the same stimulus type. This limitation prevents us from comparing the functional connectivity driven by distinct stimuli within the same category (i.e., more different clips of natural movies). On the other hand, we use a relatively wide range of natural image and natural movie stimuli, and sampled multiple stimulus types of varying complexity. While these laboratory conditions are much more controlled than natural viewing conditions, we nevertheless have determined functional connectivity under a wide range of stimulus conditions.

On the timescale of sensory processing, neuronal networks have relatively fixed anatomical connectivity. Their functional connectivity, however, can and does vary quite substantially. Our work revealed striking patterns to this functional reorganization. These patterns suggest potentially important principles governing cortical computation, such as the dynamical organization of groups of neurons into feedforward loop motifs, and the adjustment of network modularity based on stimulus complexity. Beyond their relevance for basic neuroscience, these findings may provide guidance for how to engineer dynamically robust information processing systems.

## 4 Methods

### 4.1 Dataset

We analyzed the Neuropixels dataset from Allen Institute [26]. The Neuropixels project uses high-density extracellular electrophysiology probes to record spikes from multiple regions in the mouse brain. Data used to construct functional networks are recordings of the neural activity by 6 Neuropixels probes in 6 visual cortical areas (V1, LM, RL, AL, PM, AM) from 7 mice while the mice passively viewed a visual stimulus set that contains 6 types of visual stimuli with multiple repeats: grey screen (simulation for resting-state activity), flashes, drifting gratings, static gratings, natural scenes and natural movies. On average, there are 668 ± 131 units simultaneously recorded for each mouse. In order to make a fair comparison across different visual stimuli, only neurons with a firing rate of at least 2 Hz during all stimuli are included in our analysis, thus the number of neurons (size of the functional network) is the same for each mouse given different stimuli. As a result, there are 176 ± 44 units on average for each mouse.

### 4.2 Cross-correlogram and significant functional connection

Functional connectivity is measured through Cross-correlograms (CCGs) [91]. For each stimulus type, the average CCGs across all stimulus presentations is calculated. In order to focus on the change in connectivity driven by different stimulus types, we dismissed stimulus conditions and used all presentations as trials. CCG for lagged correlation from neurons *A* to *B* is defined as

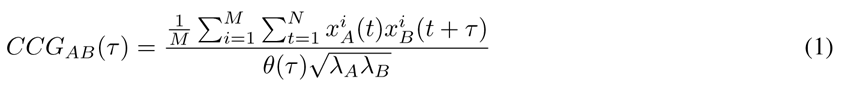

where *M* is the number of trials, *N* is the number of time bins, *x^i^* and *x^i^* are the spike trains for neuron *A* and neuron *B*, *τ* 0 is the time lag between the spike trains, *θ*(*τ*) = *M − τ* represents a triangle function that corrects for the overlap time bins, *λ_A_* and *λ_B_* are the mean firing rates for the two neurons. It is worth noting that we only allow for non-negative time lag for the sake of bidirectional connections. We used the jitter correction method to remove slow temporal correlations [92]. The jitter-corrected CCG is obtained as the difference between CCGs of the original and jittered spike trains

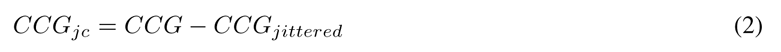

Apart from using ‘sharp peaks’ to define significant functional connections, we also included ‘sharp intervals’ to take into consideration the polysynaptic connections between neuron pairs with potentially multiple time lags. Specifically, for a given duration *D* ∈ [1, τmax + 1], where *τ_max_* = 12 ms similar to the 13 ms window in previous work [27], the set of moving average CCG is obtained by

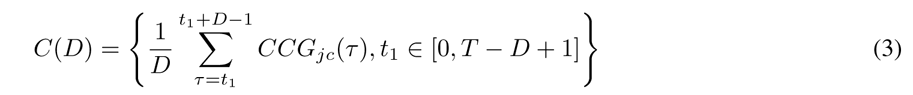

where *T* is the total length of spike trains. Therefore, there is an excitatory connection if

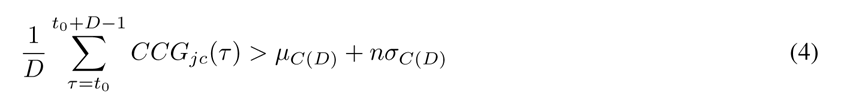

and an inhibitory connection if

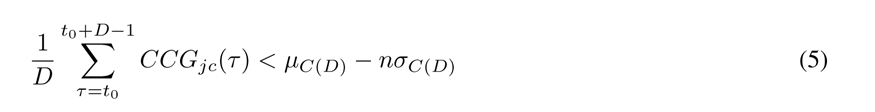

where *t*_0_ ∈ [0, τmax D + 1] is the starting time lag of the ‘sharp peak/interval’, *µ_C_*_(_*_D_*_)_*, σ_C_*_(_*_D_*_)_ are the mean and standard deviation of *C*(*D*), *n* = 4 denotes the 4-fold significance level in our experiment. It is straightforward that *D* = 1 indicates a ‘sharp peak’ while *D >* 1 denotes a ‘sharp interval’. If equation (4) or (5) is true on multiple durations *D D*_1_*, D*_2_*,, D_s_, D*_1_ *< D*_2_ *< < D_s_*, we assume the smallest duration *D* = *D*_1_ since it always leads to the highest significance level.

For bidirectional connections with zero time lag for both directions (*τ_AB_* = *τ_BA_* = 0), we only kept the direction with the higher significance level and removed the other direction unless its second highest sharp interval is also significant. Therefore, each connection was characterized by its lag, the duration of the significant interval in the CCG, and its significance value. Lag *τ* is the delay between spike trains of source neuron and target neuron, and the sign of the lag determines the direction of the connection. The duration *D* measures how long the significant peak/interval lasts, and the connection significance signals the Z score of the ‘sharp peak/interval’. Lags *τ* of across-area connections are higher than within-area connections (Supplementary Fig. 1C), which is as expected since it takes more time for a signal to travel between areas than within an area. In order to eliminate the bias brought by the lack of enough spikes or trials, we used normalized entropy for each trial to measure its statistical significance. For each neuron pair, we only keep trials in which spike trains of both neurons have a normalized entropy of at least 0.9.

### 4.3 Reference model and signed motif analysis

Since functional networks are constructed as signed networks, signed motif analysis needs to be defined. Similar to unsigned motif detection, to examine the statistical significance of signed n-neuron motifs in the networks, we generated random networks using various reference models as the baseline and conducted a comparative analysis of motif frequency between the empirical network and random networks.

Three types of commonly used reference models are adopted in this work: Erdős-Rényi model, Degree-preserving model and Pair-preserving model. However, they are all defined on unsigned networks. In order to tailor these models for analysis in the context of signed networks, we randomly assigned original edge signs to reference networks randomized using Erdős-Rényi model, Degree-preserving model and Pair-preserving model. Furthermore, we defined the Signed-pair-preserving model by preserving the edge signs for each neuron pair in the Pair-preserving model during shuffling (Supplementary Fig. 3B). Therefore, surrogate networks generated using all four reference models have the same number of positive/negative connections as the real network.

Table 1 lists the comparison between all four reference models. Erdős-Rényi model randomly shuffles connections while preserving network size, density and weight distribution [60], Degree-preserving model generates random networks while preserving size, density, weight distribution and degree distribution [93], Pair-preserving model randomizes the network while keeping size, density, weight distribution, degree distribution and neuron pair distribution [56] while the Signed-pair-preserving model preserves the signed pair distribution (Fig. 2A) in addition to the first three properties. We use Signed-pair-preserving model for signed motif analysis. For all analyses including a reference model, we randomly generated 200 surrogate networks.

**Table 1.** Reference models.

| Reference model | size & density & weight | degree distribution | pair distribution | signed pair distribution |
| --- | --- | --- | --- | --- |
| Erdős-Rényi model | ✓ | ✗ | ✗ | ✗ |
| Degree-preserving model | ✓ | ✓ | ✗ | ✗ |
| Pair-preserving model | ✓ | ✓ | ✓ | ✗ |
| Signed-pair-preserving model | ✓ | ✓ | ✓ | ✓ |

For two-neuron motif analysis, we adopted Erdős-Rényi model as the reference model and computed the relative count for each type of two-neuron connection by dividing the count of the empirical network and the average count of surrogate networks. For simplicity, we only focused on three-neuron subnetworks apart from two-neuron subnetworks during motif analysis. We used the Z score of intensity compared with reference models to determine motif significance [59]. The intensity of a certain motif *M* is defined as the summation over the intensities of all subgraphs *g* that have the structure of *M*

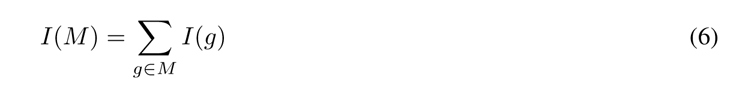

where the intensity of a certain subgraph is defined as the geometric mean of all its connection strengths

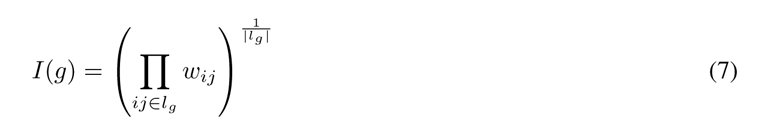

where *l_g_*denotes the set of connections in *g* and *w_ij_*is the strength of the connection from neuron *i* to *j*. Then the Z score of intensity for motif *M* can be computed as

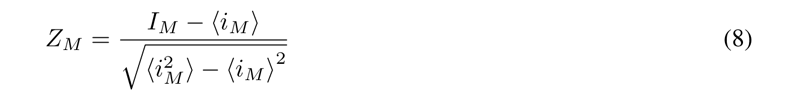

where *i_M_* is the total intensity of motif *M* in one realization of the reference model.

### 4.4 Signed module detection

The original Modularity used to detect community structure for directed networks [94] is defined as 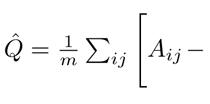 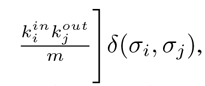, where is the adjacency matrix of the network, is the number of links, represent the in-degree and out-degree, respectively. *δ* is the Kronecker delta function and *σ_i_* denotes the community label that node *i* is assigned to. In the presence of negative links, we denote *A*^+^ = *A_ij_* if *A_ij_* ≥ 0 and zero otherwise, *A^−^* = −*A_ij_* if *A_ij_* ≤ 0 and zero otherwise, so that *A* = *A*^+^ − *A^−^*. In order to cluster nodes towards social balance, [95] proposed a frustration metric ^L^ (*λA^−^* − (1 − *λ*)*A*^+^)*δ*(*σ_i_, σ_j_*). However, neither is suitable for partitioning signed networks. In this work we adopted modified Modularity for community detection of the signed, weighted and directed CCG network. Modified Modularity of a certain partition *σ* is defined as the weighted combination of the positive and negative parts [68, 69]

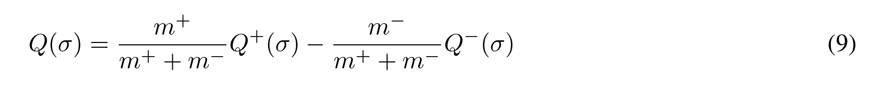

where

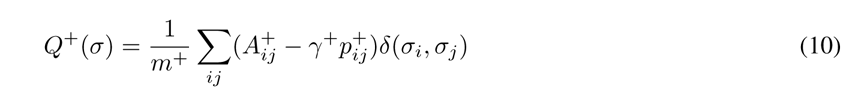

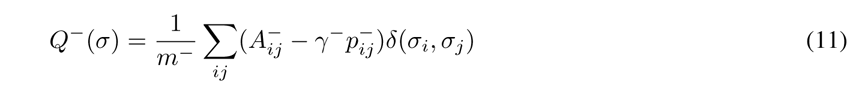

*γ*^+^ and *γ^−^* are the resolution parameters, *m*^+^ and *m^−^* are the number of positive and negative connections, respectively, *p*^+^ and *p^−^* are the connection probabilities for positive and negative links, respectively. Here we take into consideration *± out± in* degree distribution by defining the probabilities as 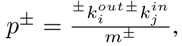, where *^±^k^out^* is the positive/negative out-degree of *_m_± i* neuron *i* and *^±^k^in^* is the positive/negative in-degree of neuron *j*. Therefore, equation (9) can be rewritten as

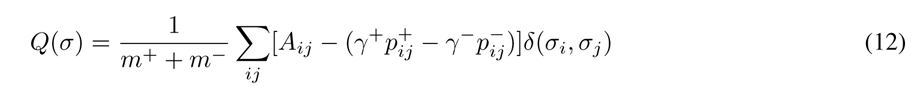

The Louvain method is a module (community) detection algorithm for partitioning networks into groups of nodes with dense connections within groups and sparse connections between groups [67]. The algorithm uses the original Modularity *Q*^^^ as a quality function to optimize the partitioning of the network . The Louvain method operates through a series of iterative steps that merge neighboring modules to maximize the Modularity gain until a locally optimal partition is reached. The algorithm uses a bottom-up approach, starting from single-node modules, and iteratively merges modules to form larger ones. To take into consideration edge signs, we revised the quality function in the Louvain method from original Modularity *Q*^^^ to modified Modularity *Q*. Therefore, the modified Louvain method aims to find an optimal partition of nodes such that positive connections are placed within modules while negative connections are between modules.

In order to determine the resolution parameters for module analysis, we obtained a Modularity difference heatmap by varying *γ*^+^ and *γ^−^* and computing the difference between Modularities of empirical and surrogate networks generated by the Signed-pair-preserving model, then looked for the *γ*^+^ and *γ^−^* that maximize the difference [14]. This way we obtained the modular partitioning that is the least random. We used the Z score of Modularity to show how modular a functional network is through comparison with a reference model (Signed-pair-preserving model). The Z score of Modularity is defined as

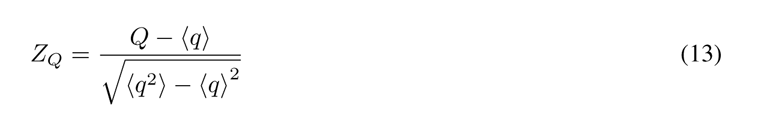

where *q* is the Modularity in one realization of the reference model. Only modules with a size of at least four neurons are included in subsequent analysis to eliminate the noise influence of isolated single neurons, pairs and triplets. Note that we included connection strength in both motif and module analyses. The CCG peak values represent connection strengths, i.e., we use absolute sum of positive/negative connection weights instead of number of positive/negative connections and the positive/negative degree of a neuron is replaced by total positive/negative connection weights. Unless otherwise stated, *Q* represents the modified Modularity for signed networks. When visualizing modular structure, the location of each node is determined by applying the Fruchterman-Reingold Layout recursively on the hypergraph and then the subgraph of each community (python package Netgraph).

### 4.5 Analysis of modular structure

To measure the fundamental properties of modular structure, we used (weighted average) coverage, (weighted average) purity and Adjusted Rand Index (ARI) to show how neurons from different visual areas are clustered together. Coverage, defined as max*_j_* 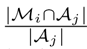 and purity, defined as max*_j_* 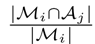, are neuron-level metrics, while their weighted averages *|Aj | |Mi|* (WA) with module size as weight are network-level metrics. The WA coverage is

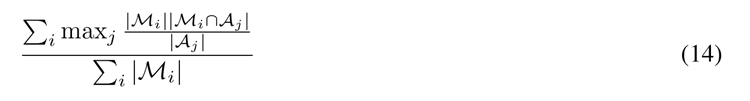

whereas the WA purity is

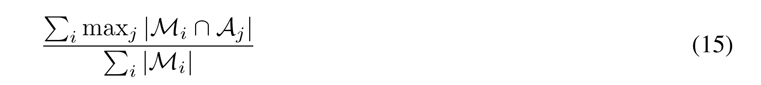

For each module partition, we also used Adjusted Rand Index (ARI) to measure its similarity to areal organization for each network. Based on the contingency table 2, ARI is defined as

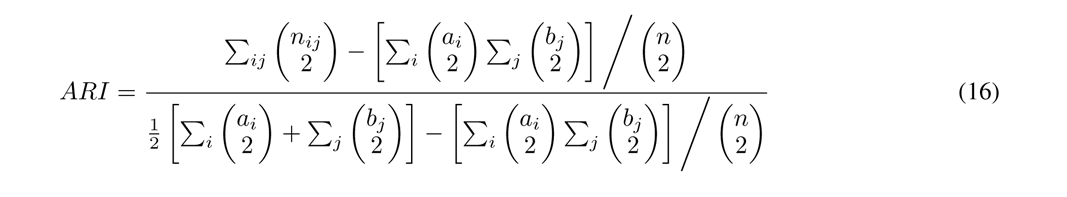

**Table 2.** Contingency table for module partition *M* = {*M*_1_, *M*_2_*, . . ., *M*_r_* and visual areal organization = {*A*_1_, *A*_2_*, . . ., *A*_s_* . *A_i_* denotes the set of neurons in the *i*-th module while *_j_* represents the set of neurons in the *Aj*-th visual area. Since we only focus on six cortical areas, thus *s* = 6. Each entry *n_ij_* denotes the number of neurons that are assigned into module M*_i_* are from visual area *A_j_*: *n_ij_* = |*M_i_* ∩ *A_j_*|.

| | $\mathcal{A}_1$ | $\mathcal{A}_2$ | ⋯ | $\mathcal{A}_s$ | sums |
| --- | --- | --- | --- | --- | --- |
| $\mathcal{M}_1$ | $n_{11}$ | $n_{12}$ | ⋯ | $n_{1s}$ | $a_1$ |
| $\mathcal{M}_2$ | $n_{21}$ | $n_{22}$ | ⋯ | $n_{2s}$ | $a_2$ |
| ⋮ | ⋮ | ⋮ | $\ddots$ | ⋮ | ⋮ |
| $\mathcal{M}_r$ | $n_{r1}$ | $n_{r2}$ | ⋯ | $n_{rs}$ | $a_r$ |
| sums | $b_1$ | $b_2$ | ⋯ | $b_s$ | |

### 4.6 Multi-resolution module partition

To reduce noise in module partition, we focused on the most active neurons that have at least 1 connection during all stimuli when examining how module partition changes with resolution parameters. Since the module partition method (modified Louvain method) is stochastic, 200 independent runs were carried out for partitioning any empirical network. To compare module partitioning results across resolution parameters, we combined multiple partitioning results based on a voting mechanism that keeps frequent modules. We first looped over each run of the modified Louvain algorithm and for each module in the run, updated the module assignment count for each node in the module. Next, we initialized a list of unassigned nodes and assigned them to the module with the highest vote count that it is partitioned into during at least one run. Each node was only assigned to one module. In each step, we removed all the nodes assigned to the most frequent module from the list of unassigned nodes and continued until all nodes were assigned to a module.

Due to the significantly greater abundance of excitatory connections compared to inhibitory connections, the parameter *γ*^+^ exerts a substantially more pronounced effect on the outcomes of module partitioning than *γ^−^*. Consequently, we limited the range of variation for *γ^−^* while placing greater emphasis on the alignment and comparison of module identity with *γ*^+^ across a broader range of values.

To compare module partitions across multiple resolutions, we assigned module IDs to modules across resolutions based on their hierarchical structure and produced a visualization of the resulting heatmap. To accomplish this, we started from the highest resolution, and traversed through the resolutions in reverse order. For the highest resolution, we assigned each module a unique ID.

Then for subsequent resolutions, we identified the largest submodule from the previous resolution and determined that its ID is inherited from previous module. To achieve this, we consider the modules from the previous resolution and calculated their overlaps with the current module. We select the submodule(s) with the maximum overlap and retrieve the corresponding ID(s) assigned to it. These ID(s) are assigned to the current module, ensuring consistency and preserving the hierarchical relationship across resolutions. For other modules that are not the largest submodule of any previous module, we assigned a new module ID to it.

Once the module IDs are assigned to the modules for all resolutions and stimuli, we sorted the nodes within each area based on their combined similarity across all stimuli to ensure an intuitive visualization. Specifically, we employed a two-opt optimization algorithm to determine an optimal node order that maximizes the similarity between module IDs of adjacent nodes across resolutions. The object is to minimize the hamming distance between 10 adjacent nodes, making neighboring nodes more likely to have similar module IDs across resolutions.

### 4.7 Statistical analysis

Since module partitioning and the generation of surrogate networks are both stochastic, each analysis involving modular structure or surrogate networks is performed with 200 independent runs. We adopt Cochran-Armitage trend test to assess the association between variables. Student’s t-test is used for the significance level between Gaussian distributions, and Shapiro-Wilk test is used for normality test. If at least one distribution is not normal, Wilcoxon rank sum test is used. Kolmogorov-Smirnov test is used to test whether a distribution is the largest among a set of distributions. Benjamini/Hochberg method is used to correct p value for false discovery rate in multiple tests.

For data with a limited number of samples, the non-parametric bootstrap method was used to calculate the confidence interval for a given sample of data. The confidence interval is based on the distribution of the medians of the bootstrap samples, which is an approximation of the sampling distribution of the median of the population from which the data were drawn. The percentile method is used to calculate the confidence interval, which involves finding the upper and lower bounds of the interval based on the percentiles of the bootstrap distribution. Specifically, we used a 95% confidence level for all the confidence intervals, the sample size was 10000.

## 5 Data availability

All the data analyzed in this manuscript is part of the Allen Brain Observatory introduced in [26]. The data used to generate main text Figs. 1–5 is available for download in Neurodata Without Borders (NWB) format via the AllenSDK. Example Jupyter Notebooks for accessing the data can be found at https://allensdk.readthedocs.io/en/latest/visual_coding_neuropixels.html.

## 6 Code availability

Code for analyses in the manuscript and generation of figures are available from the repository: https://github.com/HChoiLab/functional-network.

## 7 Acknowledgements

We thank Yu Hu, Josh Siegle, Eric Shea-Brown, and Stefan Mihalas for helpful and insightful comments on the manuscript.

## 8 Author contributions

Conceptualization (DT, JZ, XJ, HC); Methodology and Software (DT); Formal Analysis (DT); Investigation (DT, JZ, XJ, HC); Writing (DT, JZ, XJ, HC); Visualization (DT); Supervision (JZ, XJ, HC)

## 9 Funding

This work was supported by a Discovery Grant from the Natural Sciences and Engineering Research Council of Canada (RGPIN-2019-06379) and a Canada Research Chair grant to J.Z., a grant from the Tsinghua–Peking Center for Life Sciences to X.J., and the National Eye Institute of the National Institutes of Health under Award Number R00 EY030840 and a Sloan Research Fellowship to H.C. The content is solely the responsibility of the authors and does not necessarily represent the official views of the National Institutes of Health.

## 10 Competing interests

The authors declare no competing interests.

---

**Fig. S1.**
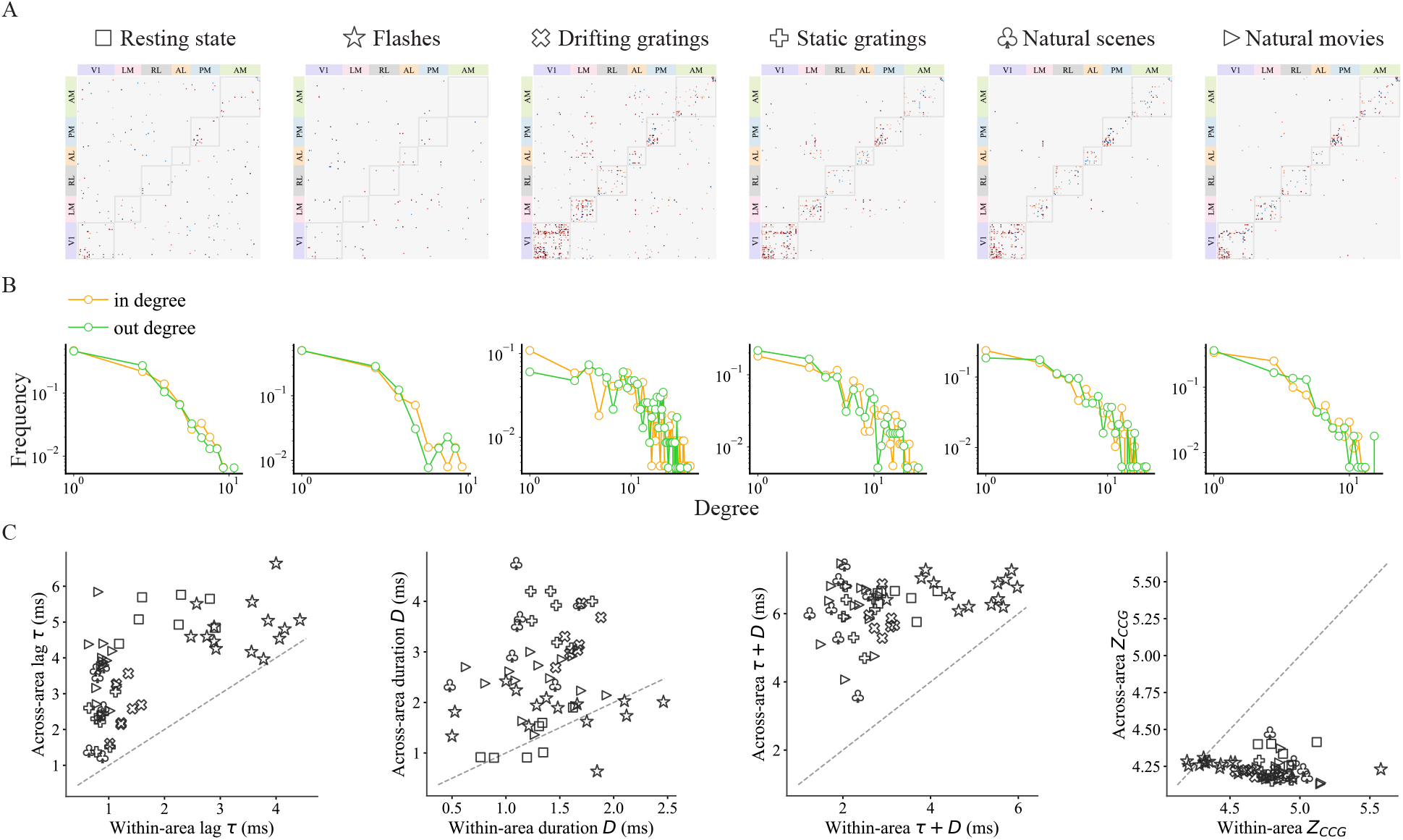
Basic properties of functional networks during all visual stimuli. (A) CCG adjacency matrices of a mouse given distinct stimuli (on the same scale as in Fig 1D). (B) Directed degree distributions of a mouse given all stimuli. (C) Across-area VS within-area comparisons for the lag *τ*, duration *D* and their sum *τ* + *D*.

**Fig. S2.**
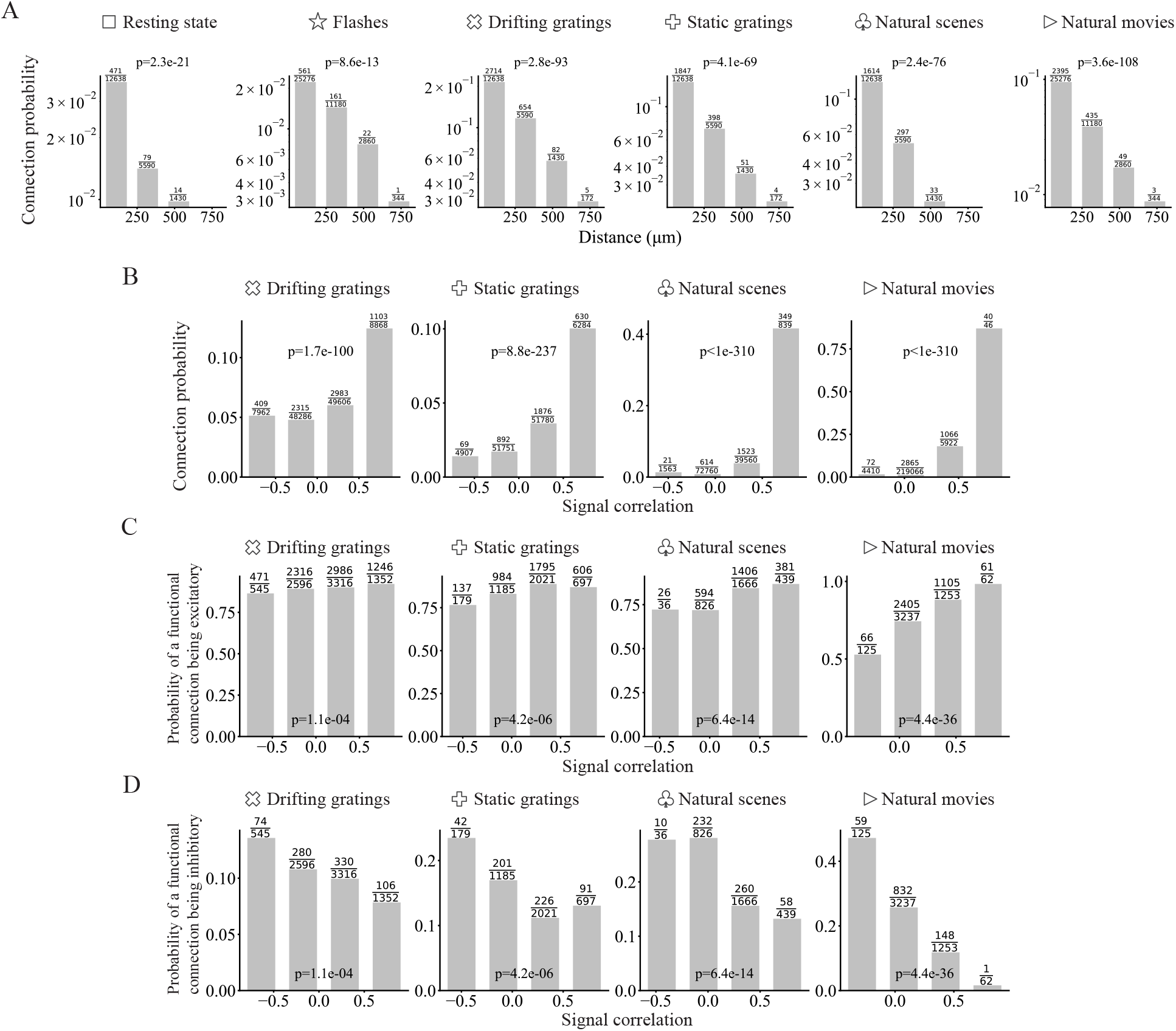
Cochran-Armitage trend test for association between (A) (functional) connection probability and distance of neurons. The rest of the plots show the Cochran-Armitage trend test for association between signal correlation and (B) (functional) connection probability, (C) probability of a functional connection being excitatory and (D) probability of a functional connection being inhibitory, under four different types of visual stimuli. Here we excluded flashes since there are only two stimulus conditions (light or dark) and signal correlation could be heavily biased and trivial.

**Fig. S3.**
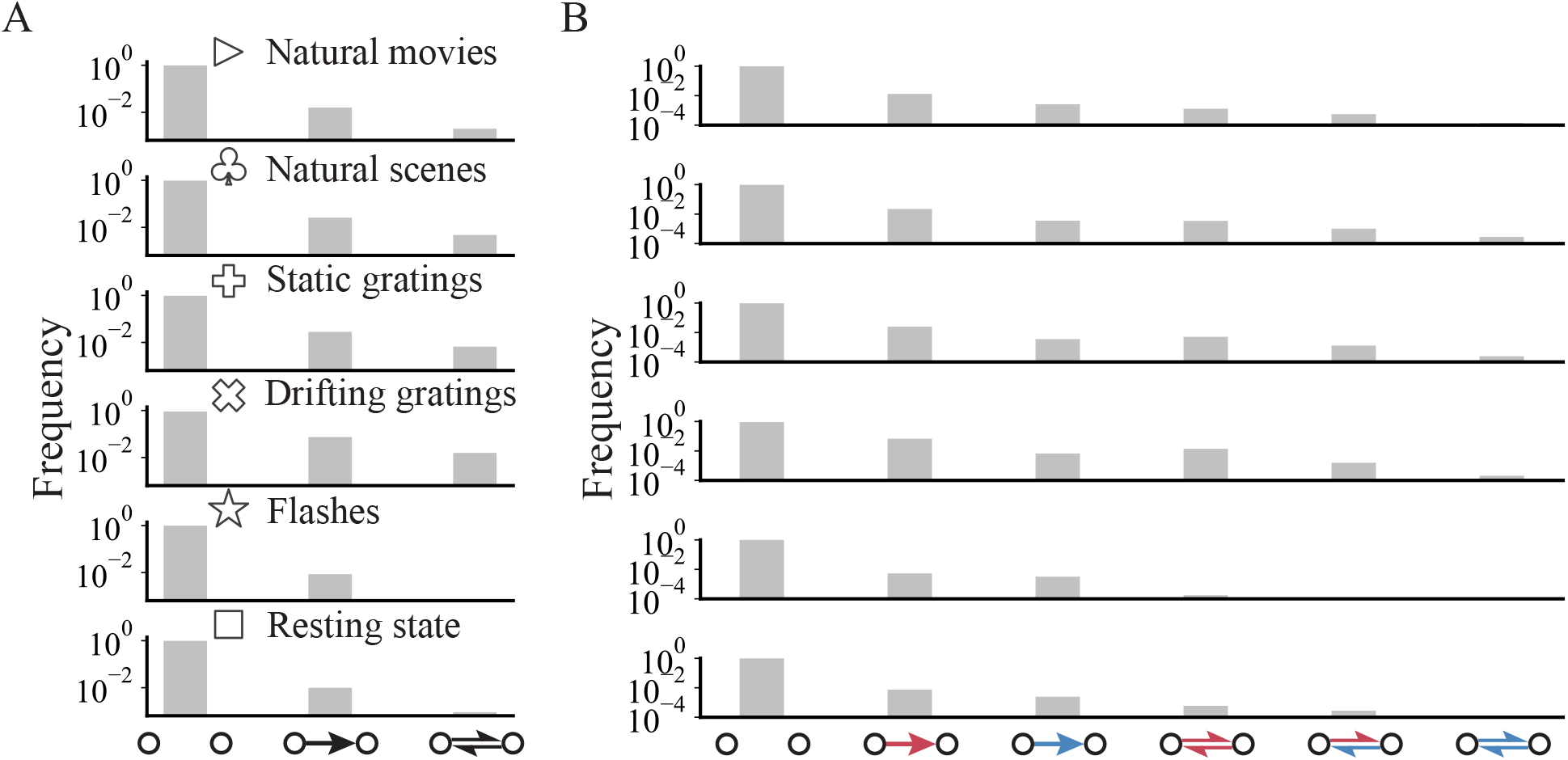
Distributions of (signed) neuron pairs. (A) Distribution of neuron pairs without edge signs. (B) Distribution of signed neuron pairs.

**Fig. S4.**
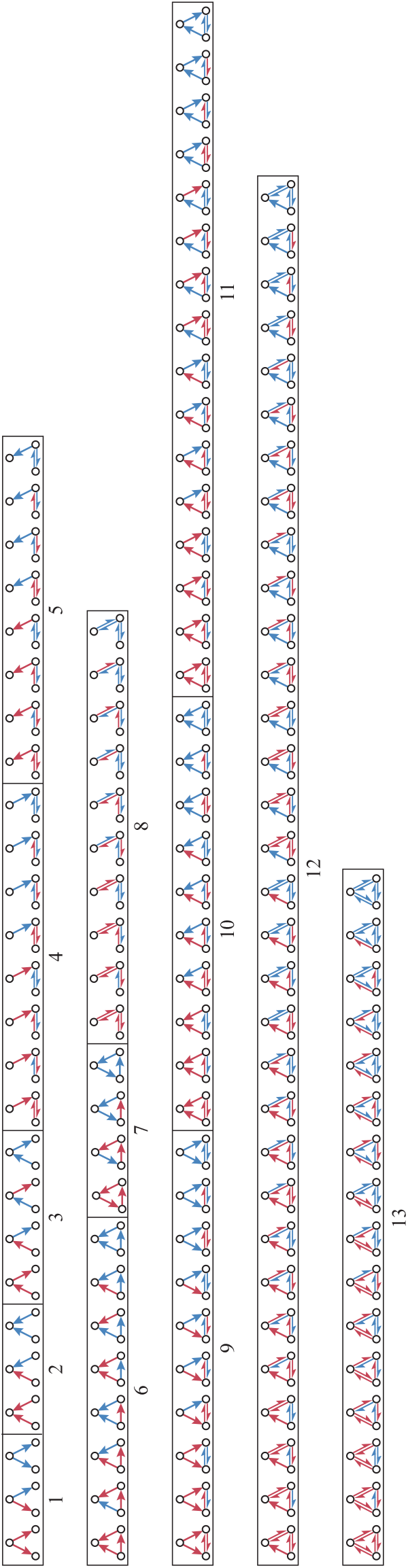
All signed motifs in the same order as Fig. 2C.

**Fig. S5.**
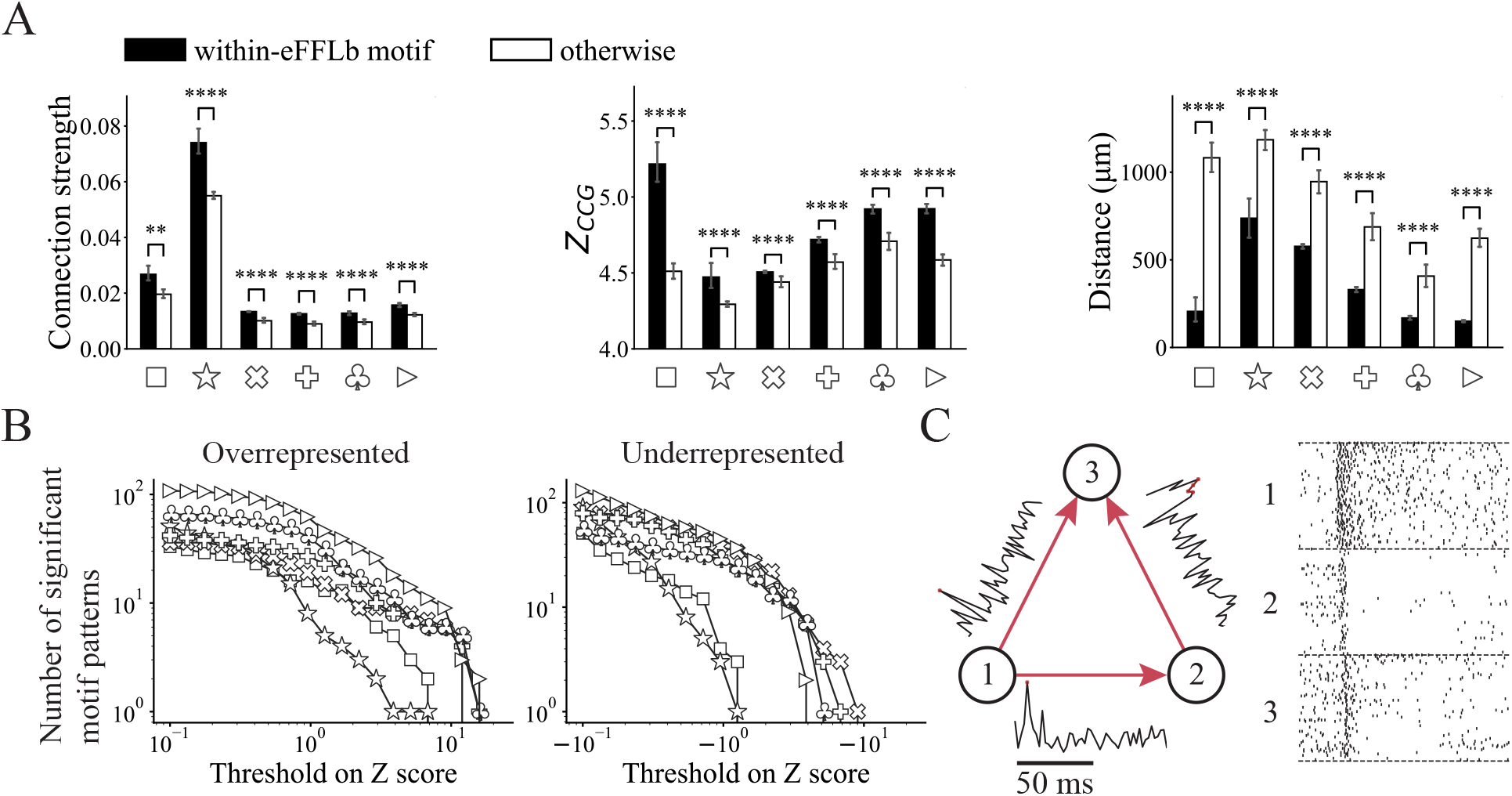
Further comparison of within-eFFLb-motif and other connections. (A) (left) Connection strength, (center) Z score of CCG and (right) physical distance for within-eFFLb-motif connections and others during all visual stimuli. *p <* 10*^−^*^2^, *p <* 10*^−^*^4^, Student’s t-test. (B) Number of significant signed motifs with intensity Z score higher than the threshold (over-represented) or lower than the threshold (under-represented) during all visual stimuli. (C) Example eFFL motif with CCGs for each connection and spike trains for each neuron.

**Fig. S6.**
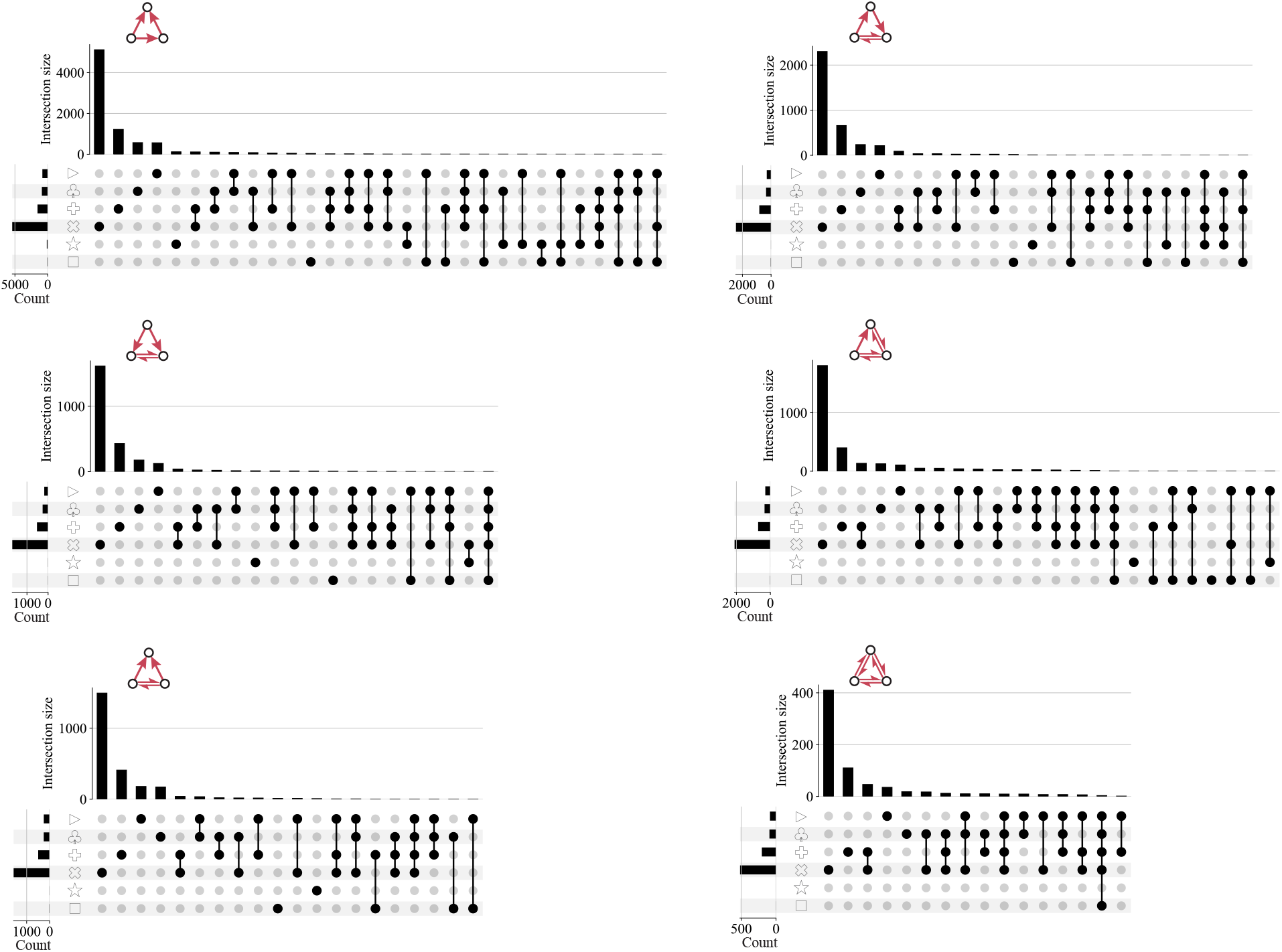
Intersections of unique motif sets for eFFLb motifs during six types of stimuli. All possible intersections with at least 1 element are shown.

**Fig. S7.**
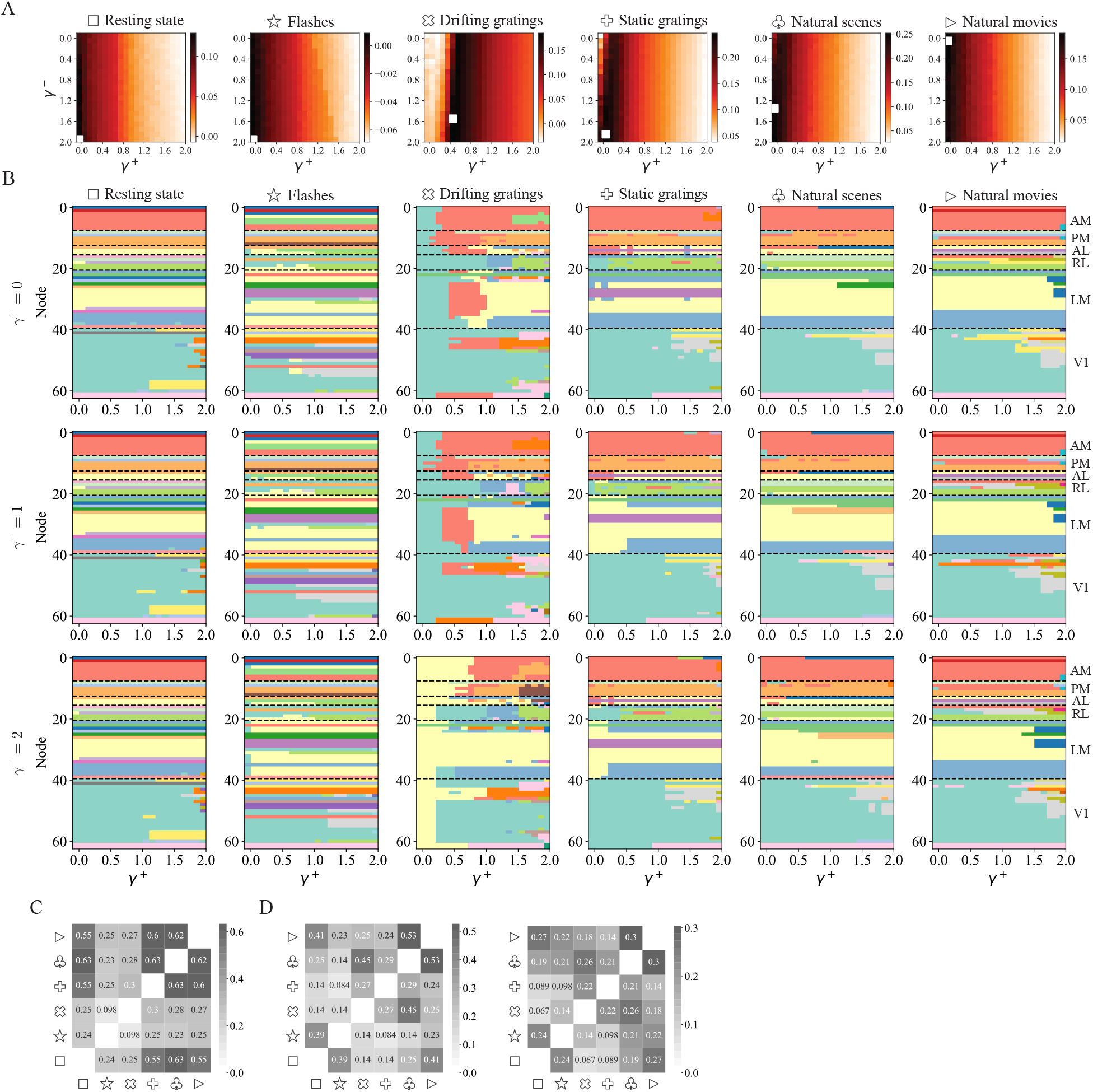
Multi-resolution modular structure. (A) The Modularity difference heatmap between the empirical network and reference model (Signed-pair-preserving model) is used to determine the resolution parameters *γ*^+^ and *γ^−^*, white box represents the maximum of the heatmap while its coordinates correspond to the optimal resolution parameters. (B) Modular partition at different resolution parameters for a mouse. Since empirical functional networks tend to have more excitatory than inhibitory connections, *γ*^+^ has a larger impact than *γ^−^* thus we only show the results obtained with three different values of *γ^−^*. Module IDs (colors) across different *γ*^+^ are determined by assigning the merged module the ID of its largest submodule at the previous step (larger resolution parameter), similar to the previous method [14]. Only neurons with at least one excitatory connection during any stimuli are included for brevity, and remaining neurons within each visual area are ordered based on their partition similarity while the module IDs across different multi-resolution modular partition maps are matched using a heuristic algorithm based on their similarity for visual comparison. (C) The heatmap of pairwise adjusted rand index (ARI) between visual stimuli for the same mouse in (B). Each multi-resolution modular partition map is considered a single clustering result, and ARI is used to measure the similarity between different partition maps. Note that ARI is independent of the color-matching heuristic algorithm and is thus more reliable. (D) The heatmaps of pairwise ARI between visual stimuli for two other mice. Only mice with at least one neuron in each area are shown. Despite individual differences, the similarity between natural scenes and movies is always among the highest.

**Fig. S8.**
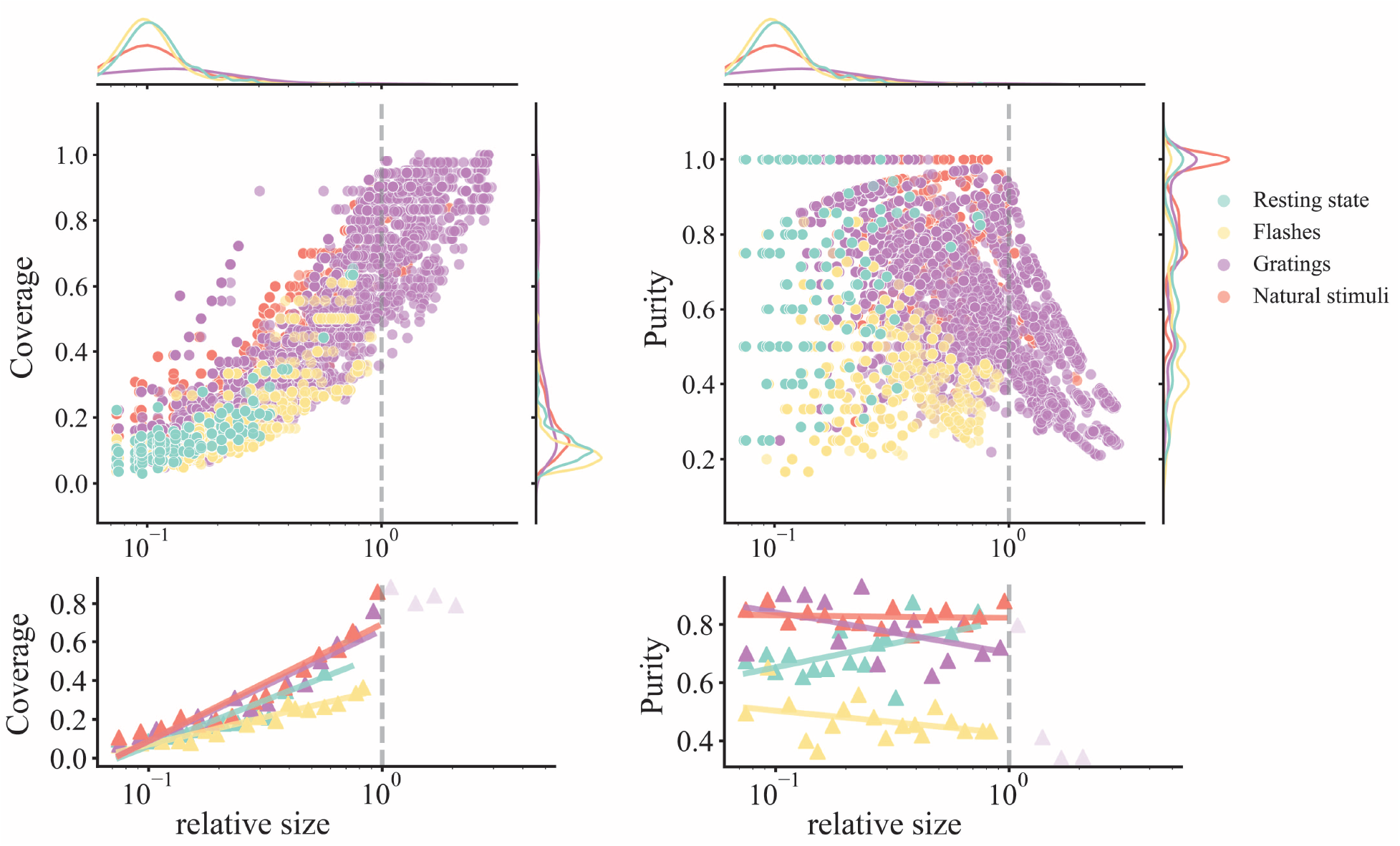
Fundamental properties of modular structure with module size. (top) Coverage/purity against relative module size for four stimulus types where each dot represents a single module; relative module size is defined as module size divided by the largest area size. (bottom) Coverage/purity against relative module size with regression after log binning. Modules with a relative size larger than 1 are excluded in regression since their coverage/purity will introduce bias.

**Fig. S9.**
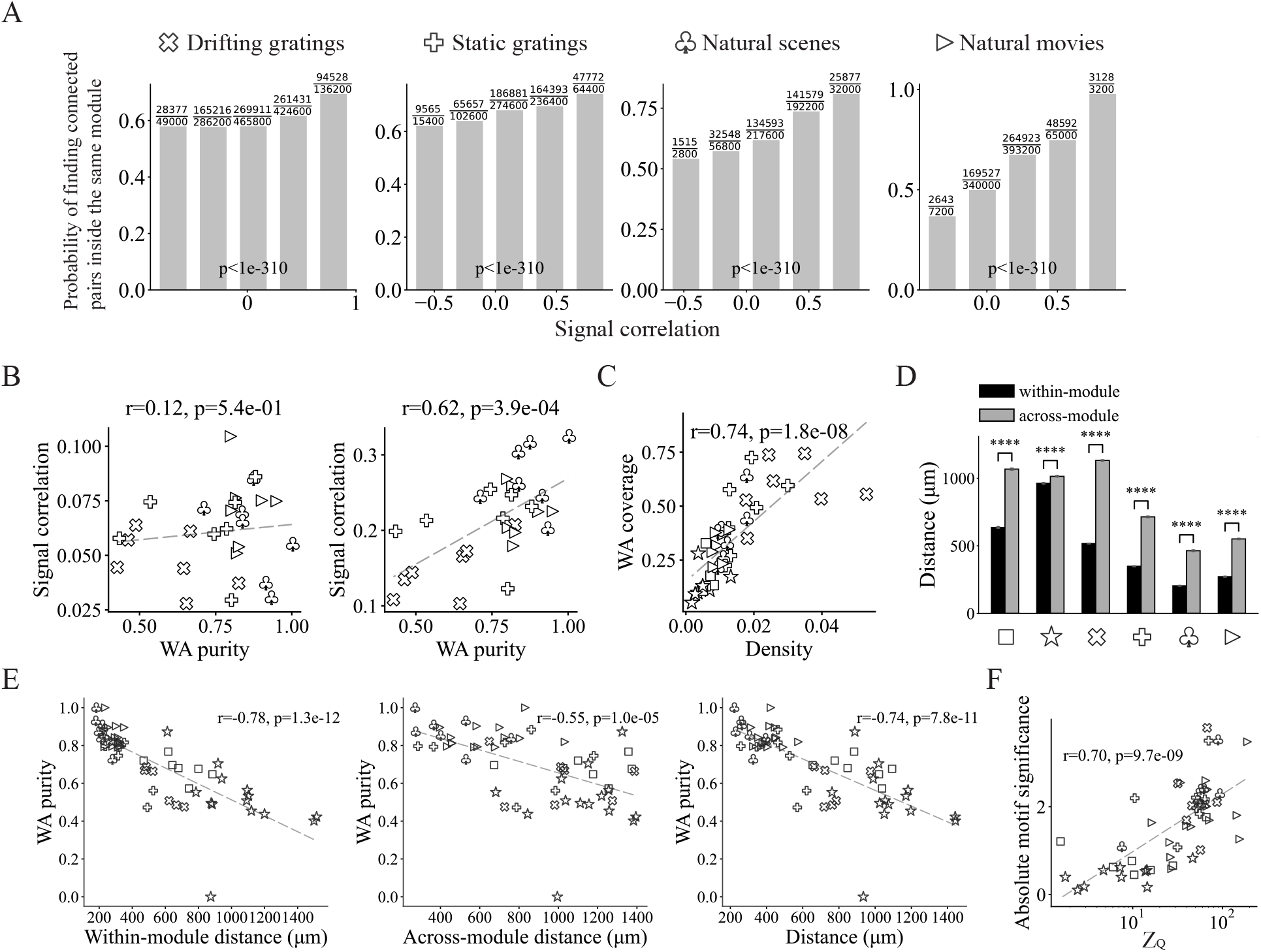
Biological interpretation of the modular structure. Wald Test is used in (B), (D) and (E). (A) Probability of finding connected neuron pairs inside the same module against their signal correlation, Cochran-Armitage trend test. (B) Linear regression results of signal correlation of chunked tuning curves with equal length against WA purity for (left) disconnected neuron pairs and (right) connected neuron pairs. Tuning curves are chunked into sequences with equal lengths for a fair comparison across stimuli, each dot represents the functional network of a mouse during certain stimulus presentations. (C) Linear regression results of WA coverage against network density. (D) Physical distance between connected neuron pairs that belong to the same or different modules. *p <* 10*^−^*^4^, Student’s t-test. (E) Linear regression results of WA purity against average distance between (left) within-module connected neuron pairs, (center) across-module connected neuron pairs and (right) all connected neuron pairs. Each dot represents the functional network of a mouse during certain stimulus presentations. (F) Average absolute motif significance (absolute Z score of intensity) across all signed motifs against Z score of Modularity.

**Fig. S10.**
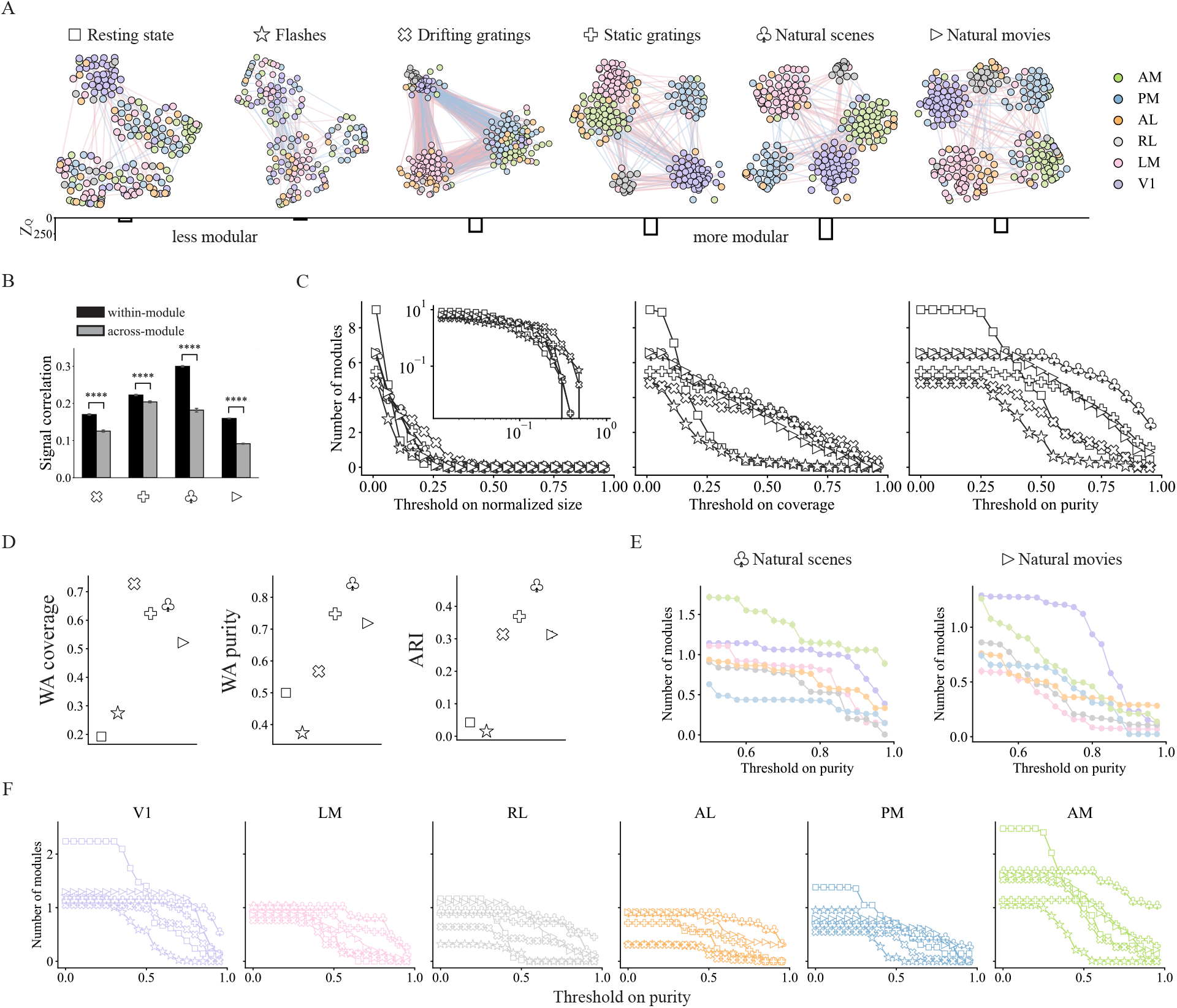
Results on modular structure with original Modularity by omitting edge signs. (A) Topological structure of functional networks during all visual stimuli. (B) Signal correlation for within-module and across-module connections. *p <* 10*^−^*^4^, Student’s t-test. (C) Number of modules with normalized size, coverage or purity higher than the threshold, inset shows the plot on a log-log scale. (D) WA (weighted average) coverage, WA purity and ARI during six visual stimuli, the error bars show the confidence intervals over all mice obtained with non-parametric bootstrap method. (E) Number of modules against threshold on purity for each visual area separately during natural scenes and natural movies. (F) Number of modules with purity higher than the threshold for each visual area during all visual stimuli.

**Fig. S11.**
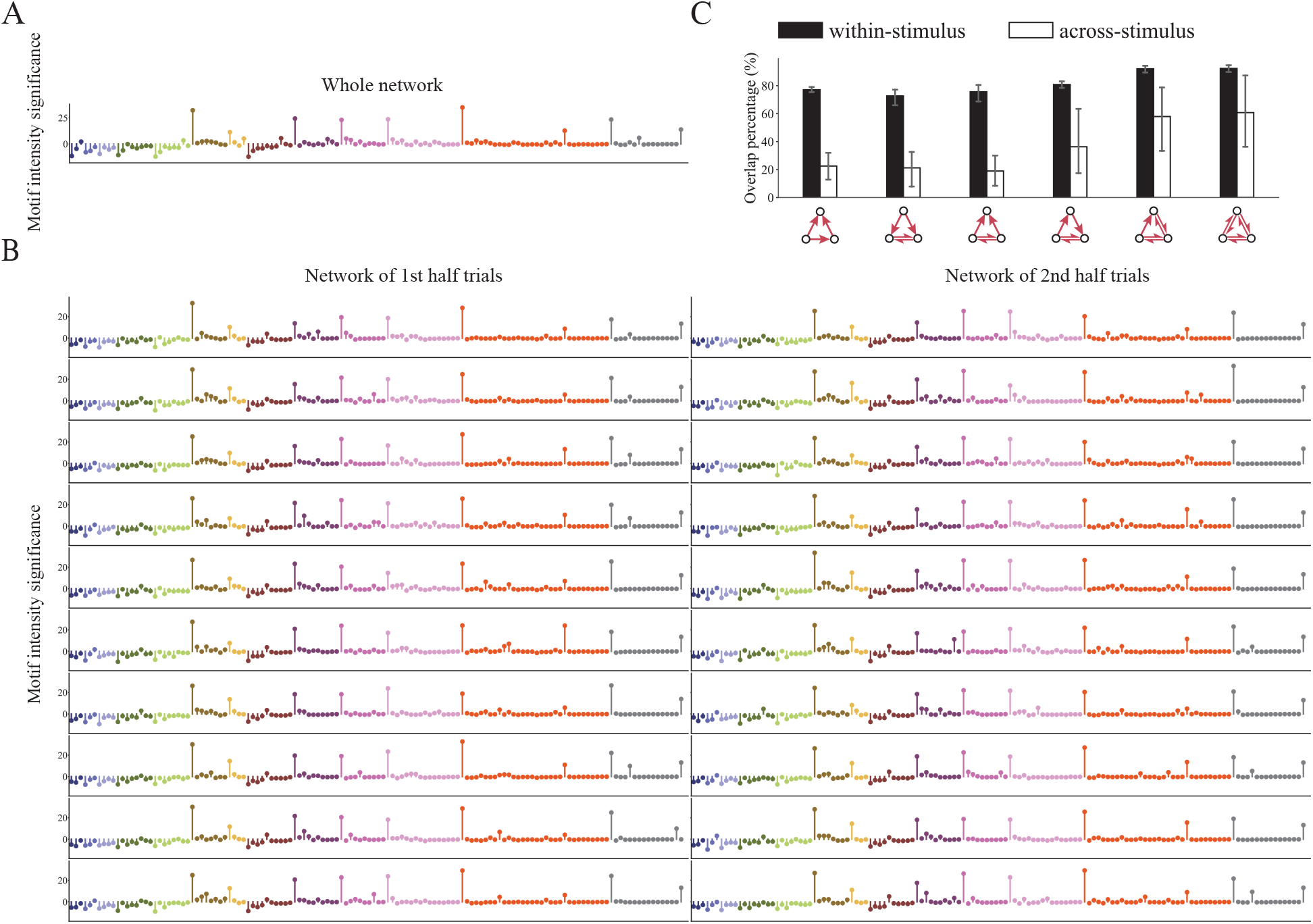
Functional networks constructed from different trials. (A) Motif sequence of the network during all trials of natural scenes. (B) Motif sequences of the networks during two halves of the trials of natural scenes. Each row corresponds to a realization of the random split. (C) Overlap percentage for eFFLb motifs. For each network, overlap percentage is defined as the percentage of motifs that are found on this network and at least one other network. Overlap percentage is higher for networks evoked by different trials from the same stimulus than different stimuli; ∗ ∗ ∗ ∗ *p <* 0.027, rank-sum t-test, corrected using Benjamini/Hochberg method.

**Fig. S12.**
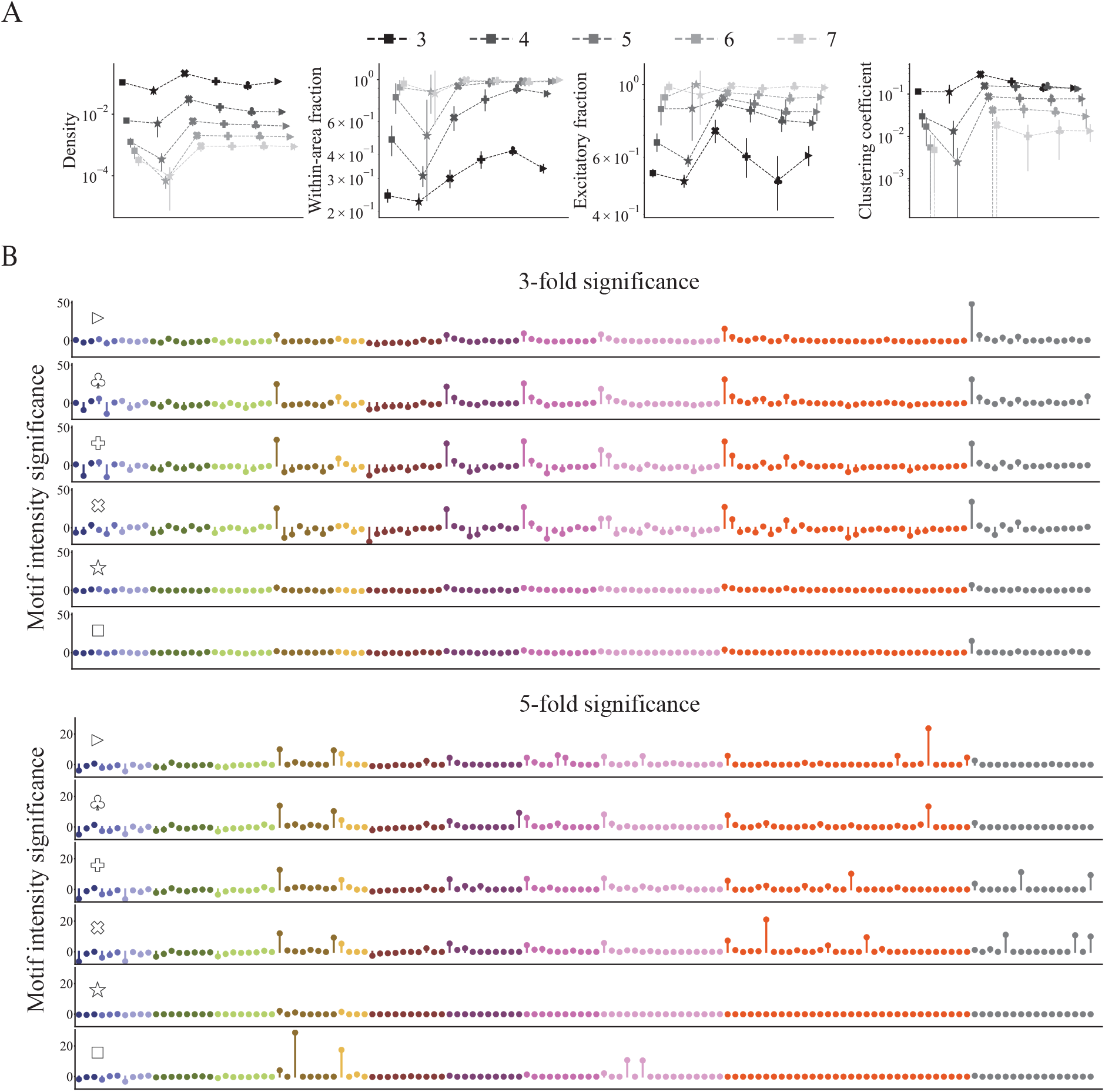
Functional networks constructed using different significance levels on functional connections. (A) Fundamental properties of the network on different significance levels (from 3-fold to 7-fold). (B) Motif sequences of the networks on different significance levels. Only 3-fold and 5-fold are shown since higher significance levels lead to extremely sparse networks. See Fig. 2C for results on 4-fold significance.

## Notes

### Competing Interest Statement

The authors have declared no competing interest.

## References

[1] Sen Song, Per Jesper Sjöström, Markus Reigl, Sacha Nelson, and Dmitri B Chklovskii. Highly nonrandom features of synaptic connectivity in local cortical circuits. PLoS Biology, 3(3):e68, 2005.

[2] Ho Ko, Sonja B Hofer, Bruno Pichler, Katherine A Buchanan, P Jesper Sjöström, and Thomas D Mrsic-Flogel. Functional specificity of local synaptic connections in neocortical networks. Nature, 473(7345):87–91, 2011.

[3] Ho Ko, Lee Cossell, Chiara Baragli, Jan Antolik, Claudia Clopath, Sonja B Hofer, and Thomas D Mrsic-Flogel. The emergence of functional microcircuits in visual cortex. Nature, 496(7443):96–100, 2013.

[4] MICrONs Consortium and, et al. Functional connectomics spanning multiple areas of mouse visual cortex. bioRxiv, 2021.

[5] Seung Wook Oh, Julie A Harris, Lydia Ng, Brent Winslow, Nicholas Cain, Stefan Mihalas, Quanxin Wang, Chris Lau, Leonard Kuan, Alex M Henry, et al. A mesoscale connectome of the mouse brain. Nature, 508(7495):207– 214, 2014.

[6] Joseph E. Knox, Kameron D. Harris, Nile Graddis, Jennifer D. Whitesell, Hongkui Zeng, Julie A. Harris, Eric Shea-Brown, and Stefan Mihalas. High-resolution data-driven model of the mouse connectome. Network Neuroscience, 3(1):217—-236, 2018.

[7] Ann M Hermundstad, Danielle S Bassett, Kevin S Brown, Elissa M Aminoff, David Clewett, Scott Freeman, Amy Frithsen, Arianne Johnson, Christine M Tipper, Michael B Miller, et al. Structural foundations of resting-state and task-based functional connectivity in the human brain. Proceedings of the National Academy of Sciences, 110(15):6169–6174, 2013.

[8] Enrique C.A. Hansen, Demian Battaglia, Andreas Spiegler, Gustavo Deco, and Viktor K. Jirsa. Functional connectivity dynamics: Modeling the switching behavior of the resting state. NeuroImage, 105(15):525–535, 2015.

[9] Mikail Rubinov and Olaf Sporns. Complex network measures of brain connectivity: uses and interpretations. NeuroImage, 52(3):1059–1069, 2010.

[10] Ed Bullmore and Olaf Sporns. Complex brain networks: graph theoretical analysis of structural and functional systems. Nature Reviews Neuroscience, 10(3):186–198, 2009.

[11] Hae-Jeong Park and Karl Friston. Structural and functional brain networks: from connections to cognition. Science, 342(6158):1238411, 2013.

[12] Valeria Della-Maggiore and Anthony R McIntosh. Time course of changes in brain activity and functional connectivity associated with long-term adaptation to a rotational transformation. Journal of Neurophysiology, 93(4):2254–2262, 2005.

[13] Casey M. Schneider-Mizell, Forrest Bodor, Agnes L. amd Collman, Derrick Brittain, Adam A. Bleckert, Sven Dorkenwald, Nicholas L. Turner, Thomas Macrina, Kisuk Lee, Ran Lu, Jingpeng Wu, and et al. Chandelier cell anatomy and function suggest a variably distributed but common signal. bioRxiv, 2020.

[14] Julie A Harris, Stefan Mihalas, Karla E Hirokawa, Jennifer D Whitesell, Hannah Choi, Amy Bernard, Phillip Bohn, Shiella Caldejon, Linzy Casal, Andrew Cho, et al. Hierarchical organization of cortical and thalamic connectivity. Nature, 575(7781):195–202, 2019.

[15] Evelyn Tang, Chad Giusti, Graham L Baum, Shi Gu, Eli Pollock, Ari E Kahn, David R Roalf, Tyler M Moore, Kosha Ruparel, Ruben C Gur, et al. Developmental increases in white matter network controllability support a growing diversity of brain dynamics. Nature Communications, 8(1):1252, 2017.

[16] Danielle S Bassett and Edward T Bullmore. Small-world brain networks revisited. The Neuroscientist, 23(5):499– 516, 2017.

[17] Hannah Choi and Stefan Mihalas. Synchronization dependent on spatial structures of a mesoscopic whole-brain network. PLOS Computational Biology, 15(4):e1006978, 2019.

[18] Christopher J Honey, Rolf Kötter, Michael Breakspear, and Olaf Sporns. Network structure of cerebral cortex shapes functional connectivity on multiple time scales. Proceedings of the National Academy of Sciences, 104(24):10240–10245, 2007.

[19] Christopher J Honey, Olaf Sporns, Leila Cammoun, Xavier Gigandet, Jean-Philippe Thiran, Reto Meuli, and Patric Hagmann. Predicting human resting-state functional connectivity from structural connectivity. Proceedings of the National Academy of Sciences, 106(6):2035–2040, 2009.

[20] Zhuokun Ding, Paul G. Fahey, Stelios Papadopoulous, Eric Wang, Brendan Celli, and, et al. Functional connectomics reveals general wiring rule in mouse visual cortex. bioRxiv, 2023.

[21] Adam Kohn and Matthew A Smith. Stimulus dependence of neuronal correlation in primary visual cortex of the macaque. Journal of Neuroscience, 25(14):3661–3673, 2005.

[22] Kevan AC Martin and Sylvia Schröder. Functional heterogeneity in neighboring neurons of cat primary visual cortex in response to both artificial and natural stimuli. Journal of Neuroscience, 33(17):7325–7344, 2013.

[23] Pietro Berkes, Gergő Orbán, Máté Lengyel, and József Fiser. Spontaneous cortical activity reveals hallmarks of an optimal internal model of the environment. Science, 331(6013):83–87, 2011.

[24] Michael Okun, Nicholas A Steinmetz, Lee Cossell, M Florencia Iacaruso, Ho Ko, Péter Barthó, Tirin Moore, Sonja B Hofer, Thomas D Mrsic-Flogel, Matteo Carandini, et al. Diverse coupling of neurons to populations in sensory cortex. Nature, 521(7553):511–515, 2015.

[25] Mihály Bányai, Andreea Lazar, Liane Klein, Johanna Klon-Lipok, Marcell Stippinger, Wolf Singer, and Gergő Orbán. Stimulus complexity shapes response correlations in primary visual cortex. Proceedings of the National Academy of Sciences, 116(7):2723–2732, 2019.

[26] Joshua H Siegle, Xiaoxuan Jia, Séverine Durand, Sam Gale, Corbett Bennett, Nile Graddis, Greggory Heller, Tamina K Ramirez, Hannah Choi, Jennifer A Luviano, et al. Survey of spiking in the mouse visual system reveals functional hierarchy. Nature, 592(7852):86–92, 2021.

[27] Xiaoxuan Jia, Joshua H Siegle, Séverine Durand, Greggory Heller, Tamina K Ramirez, Christof Koch, and Shawn R Olsen. Multi-regional module-based signal transmission in mouse visual cortex. Neuron, 110(9):1585– 1598, 2022.

[28] Laurel Joy Gabard-Durnam, Dylan Grace Gee, Bonnie Goff, Jessica Flannery, Eva Telzer, Kathryn Leigh Humphreys, Daniel Stephen Lumian, Dominic Stephen Fareri, Christina Caldera, and Nim Tottenham. Stimulus-elicited connectivity influences resting-state connectivity years later in human development: a prospective study. Journal of Neuroscience, 36(17):4771–4784, 2016.

[29] Vincent G van de Ven, Elia Formisano, David Prvulovic, Christian H Roeder, and David EJ Linden. Functional connectivity as revealed by spatial independent component analysis of fMRI measurements during rest. Human Brain Mapping, 22(3):165–178, 2004.

[30] Fenna M Krienen, BT Thomas Yeo, and Randy L Buckner. Reconfigurable task-dependent functional coupling modes cluster around a core functional architecture. Philosophical Transactions of the Royal Society B: Biological Sciences, 369(1653):20130526, 2014.

[31] Erhan Genç, Marieke Louise Schölvinck, Johanna Bergmann, Wolf Singer, and Axel Kohler. Functional connectivity patterns of visual cortex reflect its anatomical organization. Cerebral Cortex, 26(9):3719–3731, 2016.

[32] Felice T Sun, Lee M Miller, and Mark D’esposito. Measuring interregional functional connectivity using coherence and partial coherence analyses of fMRI data. NeuroImage, 21(2):647–658, 2004.

[33] Baxter P Rogers, Victoria L Morgan, Allen T Newton, and John C Gore. Assessing functional connectivity in the human brain by fmri. Magnetic Resonance Imaging, 25(10):1347–1357, 2007.

[34] Michael W Cole, Danielle S Bassett, Jonathan D Power, Todd S Braver, and Steven E Petersen. Intrinsic and task-evoked network architectures of the human brain. Neuron, 83(1):238–251, 2014.

[35] Javier Gonzalez-Castillo and Peter A Bandettini. Task-based dynamic functional connectivity: Recent findings and open questions. NeuroImage, 180:526–533, 2018.

[36] Lauren K Lynch, Kun-Han Lu, Haiguang Wen, Yizhen Zhang, Andrew J Saykin, and Zhongming Liu. Task-evoked functional connectivity does not explain functional connectivity differences between rest and task conditions. Human Brain Mapping, 39(12):4939–4948, 2018.

[37] Axel Frien and Reinhard Eckhorn. Functional coupling shows stronger stimulus dependency for fast oscillations than for low-frequency components in striate cortex of awake monkey. European Journal of Neuroscience, 12(4):1466–1478, 2000.

[38] Sonja B Hofer, Ho Ko, Bruno Pichler, Joshua Vogelstein, Hana Ros, Hongkui Zeng, Ed Lein, Nicholas A Lesica, and Thomas D Mrsic-Flogel. Differential connectivity and response dynamics of excitatory and inhibitory neurons in visual cortex. Nature Neuroscience, 14(8):1045–1052, 2011.

[39] Ian Nauhaus, Laura Busse, Matteo Carandini, and Dario L Ringach. Stimulus contrast modulates functional connectivity in visual cortex. Nature Neuroscience, 12(1):70–76, 2009.

[40] Carsen Stringer, Marius Pachitariu, Nicholas Steinmetz, Matteo Carandini, and Kenneth D Harris. High-dimensional geometry of population responses in visual cortex. Nature, 571(7765):361–365, 2019.

[41] Nadav Kashtan, Shalev Itzkovitz, Ron Milo, and Uri Alon. Topological generalizations of network motifs. Physical Review E, 70(3):031909, 2004.

[42] Shuzo Sakata, Yusuke Komatsu, and Tetsuo Yamamori. Local design principles of mammalian cortical networks. Neuroscience Research, 51(3):309–315, 2005.

[43] Olaf Sporns, Giulio Tononi, and Gerald M Edelman. Theoretical neuroanatomy: relating anatomical and functional connectivity in graphs and cortical connection matrices. Cerebral Cortex, 10(2):127–141, 2000.

[44] Lazaros K Gallos, Hernán A Makse, and Mariano Sigman. A small world of weak ties provides optimal global integration of self-similar modules in functional brain networks. Proceedings of the National Academy of Sciences, 109(8):2825–2830, 2012.

[45] MS Berry and VW Pentreath. Criteria for distinguishing between monosynaptic and polysynaptic transmission. Brain Research, 105(1):1–20, 1976.

[46] R Clay Reid and Jose-Manuel Alonso. Specificity of monosynaptic connections from thalamus to visual cortex. Nature, 378(6554):281–284, 1995.

[47] Harvey A Swadlow and Jose-Manuel Alonso. Multielectrodes join the connectome. Nature Methods, 14(9):847– 848, 2017.

[48] Vito Paolo Pastore, Paolo Massobrio, Aleksandar Godjoski, and Sergio Martinoia. Identification of excitatory-inhibitory links and network topology in large-scale neuronal assemblies from multi-electrode recordings. PLoS Computational Biology, 14(8):e1006381, 2018.

[49] Albert-László Barabási and Réka Albert. Emergence of scaling in random networks. Science, 286(5439):509–512, 1999.

[50] Martijn P Van Den Heuvel and Hilleke E Hulshoff Pol. Exploring the brain network: a review on resting-state fmri functional connectivity. European Neuropsychopharmacology, 20(8):519–534, 2010.

[51] Cornelis Jan Stam, Willem De Haan, ABFJ Daffertshofer, BF Jones, I Manshanden, Anne-Marie van Cappellen van Walsum, Teresa Montez, JPA Verbunt, Jan C De Munck, Bob Wilhelm Van Dijk, et al. Graph theoretical analysis of magnetoencephalographic functional connectivity in Alzheimer’s disease. Brain, 132(1):213–224, 2009.

[52] Alexander S Ecker, Philipp Berens, Georgios A Keliris, Matthias Bethge, Nikos K Logothetis, and Andreas S Tolias. Decorrelated neuronal firing in cortical microcircuits. Science, 327(5965):584–587, 2010.

[53] William E Vinje and Jack L Gallant. Sparse coding and decorrelation in primary visual cortex during natural vision. Science, 287(5456):1273–1276, 2000.

[54] Rodrigo Perin, Thomas K Berger, and Henry Markram. A synaptic organizing principle for cortical neuronal groups. Proceedings of the National Academy of Sciences, 108(13):5419–5424, 2011.

[55] Bruno B Averbeck, Peter E Latham, and Alexandre Pouget. Neural correlations, population coding and computation. Nature Reviews Neuroscience, 7(5):358–366, 2006.

[56] Ron Milo, Shai Shen-Orr, Shalev Itzkovitz, Nadav Kashtan, Dmitri Chklovskii, and Uri Alon. Network motifs: simple building blocks of complex networks. Science, 298(5594):824–827, 2002.

[57] Xiaolong Jiang, Shan Shen, Cathryn R Cadwell, Philipp Berens, Fabian Sinz, Alexander S Ecker, Saumil Patel, and Andreas S Tolias. Principles of connectivity among morphologically defined cell types in adult neocortex. Science, 350(6264):aac9462, 2015.

[58] Olaf Sporns and Rolf Kötter. Motifs in brain networks. PLoS Biology, 2(11):e369, 2004.

[59] Jukka-Pekka Onnela, Jari Saramäki, János Kertész, and Kimmo Kaski. Intensity and coherence of motifs in weighted complex networks. Physical Review E, 71(6):065103, 2005.

[60] Paul Erdős, Alfréd Rényi, et al. On the evolution of random graphs. Publication of the Mathematical Institute of the Hungarian Academy of Sciences, 5(1):17–60, 1960.

[61] Uri Alon. Network motifs: theory and experimental approaches. Nature Reviews Genetics, 8(6):450–461, 2007.

[62] Chunguang Li. Functions of neuronal network motifs. Physical Review E, 78(3):037101, 2008.

[63] Olaf Sporns. Contributions and challenges for network models in cognitive neuroscience. Nature Neuroscience, 17(5):652–660, 2014.

[64] Alex Fornito, Andrew Zalesky, and Edward Bullmore. Fundamentals of brain network analysis. Academic Press, 2016.

[65] Olaf Sporns and Richard F Betzel. Modular brain networks. Annual Review of Psychology, 67:613, 2016.

[66] Ed Bullmore and Olaf Sporns. The economy of brain network organization. Nature Reviews Neuroscience, 13(5):336–349, 2012.

[67] Vincent D Blondel, Jean-Loup Guillaume, Renaud Lambiotte, and Etienne Lefebvre. Fast unfolding of communities in large networks. Journal of Statistical Mechanics: Theory and Experiment, 2008(10):P10008, 2008.

[68] Vincent A Traag and Jeroen Bruggeman. Community detection in networks with positive and negative links. Physical Review E, 80(3):036115, 2009.

[69] Sergio Gómez, Pablo Jensen, and Alex Arenas. Analysis of community structure in networks of correlated data. Physical Review E, 80(1):016114, 2009.

[70] Vincent A Traag, Gautier Krings, and Paul Van Dooren. Significant scales in community structure. Scientific Reports, 3(1):1–10, 2013.

[71] Richard F Betzel, Alessandra Griffa, Andrea Avena-Koenigsberger, Joaquín Goñi, Jean-Philippe Thiran, Patric Hagmann, and Olaf Sporns. Multi-scale community organization of the human structural connectome and its relationship with resting-state functional connectivity. Network Science, 1(3):353–373, 2013.

[72] Chunshui Yu, Yuan Zhou, Yong Liu, Tianzi Jiang, Haiwei Dong, Yunting Zhang, and Martin Walter. Functional segregation of the human cingulate cortex is confirmed by functional connectivity based neuroanatomical parcellation. NeuroImage, 54(4):2571–2581, 2011.

[73] Matthias Beckmann, Heidi Johansen-Berg, and Matthew FS Rushworth. Connectivity-based parcellation of human cingulate cortex and its relation to functional specialization. Journal of Neuroscience, 29(4):1175–1190, 2009.

[74] Rogier B Mars, Saad Jbabdi, Jérôme Sallet, Jill X O’Reilly, Paula L Croxson, Etienne Olivier, MaryAnn P Noonan, Caroline Bergmann, Anna S Mitchell, Mark G Baxter, et al. Diffusion-weighted imaging tractography-based parcellation of the human parietal cortex and comparison with human and macaque resting-state functional connectivity. Journal of Neuroscience, 31(11):4087–4100, 2011.

[75] Nadav Kashtan and Uri Alon. Spontaneous evolution of modularity and network motifs. Proceedings of the National Academy of Sciences, 102(39):13773–13778, 2005.

[76] Yu Hu, James Trousdale, Krešimir Josić, and Eric Shea-Brown. Motif statistics and spike correlations in neuronal networks. Journal of Statistical Mechanics: Theory and Experiment, 2013(03):P03012, 2013.

[77] Yu Hu, James Trousdale, Krešimir Josić, and Eric Shea-Brown. Local paths to global coherence: Cutting networks down to size. Physical Review E, 89(3):032802, 2014.

[78] Gabriel Koch Ocker, Ashok Litwin-Kumar, and Brent Doiron. Self-organization of microcircuits in networks of spiking neurons with plastic synapses. PLoS Computational Biology, 11(8):e1004458, 2015.

[79] Yu Hu, Steven L Brunton, Nicholas Cain, Stefan Mihalas, J Nathan Kutz, and Eric Shea-Brown. Feedback through graph motifs relates structure and function in complex networks. Physical Review E, 98(6):062312, 2018.

[80] Kyle Bojanek, Yuqing Zhu, and Jason MacLean. Cyclic transitions between higher order motifs underlie sustained asynchronous spiking in sparse recurrent networks. PLoS Computational Biology, 16(9):e1007409, 2020.

[81] Thomas E Gorochowski, Claire S Grierson, and Mario Di Bernardo. Organization of feed-forward loop motifs reveals architectural principles in natural and engineered networks. Science Advances, 4(3):eaap9751, 2018.

[82] Gustavo Deco and Morten L Kringelbach. Metastability and coherence: extending the communication through coherence hypothesis using a whole-brain computational perspective. Trends in Neurosciences, 39(3):125–135, 2016.

[83] R Matthew Hutchison, Thilo Womelsdorf, Elena A Allen, Peter A Bandettini, Vince D Calhoun, Maurizio Corbetta, Stefania Della Penna, Jeff H Duyn, Gary H Glover, Javier Gonzalez-Castillo, et al. Dynamic functional connectivity: promise, issues, and interpretations. NeuroImage, 80:360–378, 2013.

[84] Olaf Sporns, Giulio Tononi, and Gerald M Edelman. Connectivity and complexity: the relationship between neuroanatomy and brain dynamics. Neural Networks, 13(8-9):909–922, 2000.

[85] Valerio Mante, Vincent Bonin, and Matteo Carandini. Functional mechanisms shaping lateral geniculate responses to artificial and natural stimuli. Neuron, 58(4):625–638, 2008.

[86] Sarah Feldt, Paolo Bonifazi, and Rosa Cossart. Dissecting functional connectivity of neuronal microcircuits: experimental and theoretical insights. Trends in Neurosciences, 34(5):225–236, 2011.

[87] Yael Adini, Dov Sagi, and Misha Tsodyks. Excitatory–inhibitory network in the visual cortex: Psychophysical evidence. Proceedings of the National Academy of Sciences, 94(19):10426–10431, 1997.

[88] David Ferster and Kenneth D Miller. Neural mechanisms of orientation selectivity in the visual cortex. Annual Review of Neuroscience, 23(1):441–471, 2000.

[89] Marius Pachitariu, Nicholas Steinmetz, Shabnam Kadir, Matteo Carandini, and Harris Kenneth D. Kilosort: realtime spike-sorting for extracellular electrophysiology with hundreds of channels. bioRxiv, page 061481, 2016.

[90] Alessio P Buccino, Cole L Hurwitz, Samuel Garcia, Jeremy Magland, Joshua H Siegle, Roger Hurwitz, and Matthias H Hennig. Spikeinterface, a unified framework for spike sorting. Elife, 9:e61834, 2020.

[91] Matthew A Smith and Adam Kohn. Spatial and temporal scales of neuronal correlation in primary visual cortex. Journal of Neuroscience, 28(48):12591–12603, 2008.

[92] Matthew T Harrison and Stuart Geman. A rate and history-preserving resampling algorithm for neural spike trains. Neural Computation, 21(5):1244–1258, 2009.

[93] Sergei Maslov and Kim Sneppen. Specificity and stability in topology of protein networks. Science, 296(5569):910– 913, 2002.

[94] Elizabeth A Leicht and Mark EJ Newman. Community structure in directed networks. Physical Review Letters, 100(11):118703, 2008.

[95] Patrick Doreian and Andrej Mrvar. A partitioning approach to structural balance. Social Networks, 18(2):149–168, 1996.

